# Inhibition and Crystal Structure of the Human DHTKD1-Thiamin Diphosphate Complex

**DOI:** 10.1101/2020.01.20.913012

**Authors:** João Leandro, Susmita Khamrui, Hui Wang, Chalada Suebsuwong, Natalia S. Nemeria, Khoi Huynh, Moses Moustakim, Cody Secor, May Wang, Tetyana Dodatko, Brandon Stauffer, Christopher G. Wilson, Chunli Yu, Michelle R. Arkin, Frank Jordan, Roberto Sanchez, Robert J. DeVita, Michael B. Lazarus, Sander M. Houten

## Abstract

DHTKD1 is the E1 component of the 2-oxoadipic acid dehydrogenase complex (OADHc), which functions in the L-lysine degradation pathway. Mutations in *DHTKD1* have been associated with 2-aminoadipic and 2-oxoadipic aciduria, Charcot-Marie-Tooth disease type 2Q (CMT2Q) and eosinophilic esophagitis (EoE). A crystal structure and inhibitors of DHTKD1 could improve the understanding of these clinically distinct disorders, but are currently not available. Here we report the identification of adipoylphosphonic acid and tenatoprazole as DHTKD1 inhibitors using targeted and high throughput screening, respectively. We furthermore elucidate the DHTKD1 crystal structure with thiamin diphosphate bound at 2.1 Å. The protein assembles as a dimer with residues from both monomers contributing to cofactor binding. We also report the impact of ten *DHTKD1* missense mutations on the encoded proteins by enzyme kinetics, thermal stability and structural modeling. Some DHTKD1 variants displayed impaired folding (S777P and S862I), whereas other substitutions rendered the enzyme inactive (L234G, R715C and R455Q) or affected the thermal stability and catalytic efficiency (V360A and P773L). Three variants (R163Q, Q305H and G729R) surprisingly showed wild type like properties. Our work provides a structural basis for further understanding of the function of DHTKD1 and a starting point for selective small molecule inhibitors of the enzyme, which could help tease apart the role of this enzyme in several human pathologies.

## Introduction

DHTKD1 is the E1 component (E1a) of the 2-oxoadipate dehydrogenase complex (OADHc) that operates in L-lysine, L-hydroxylysine and L-tryptophan degradation and converts 2-oxoadipic acid (OA) into glutaryl-CoA (Figure S1) (1–7). DHTKD1 is a homolog of 2-oxoglutarate dehydrogenase OGDH (∼39% identity), the E1 component (E1o) of the 2-oxoglutarate dehydrogenase complex (OGDHc) that functions in the citric acid cycle. In general, 2-oxo acid dehydrogenase complexes have unique E1 and E2 components, but share the same E3 component. DHTKD1 and OGDH are an exception, because they share both the E2 (DLST) and E3 (DLD) components (4, 8), which allows for functional and regulatory cross-talk between the citric acid cycle and L-lysine degradation (5, 8).

Genetic studies have implicated DHTKD1 in the pathogenesis of several human diseases. Autosomal recessive mutants in *DHTKD1* cause 2-aminoadipic and 2-oxoadipic aciduria (AMOXAD; MIM 204750) (2, 3, 6), a biochemical abnormality of questionable clinical significance that is characterized by the accumulation of 2-aminoadipic acid (AAA) and OA (Figure S1) (9). Autosomal dominant variants have been associated with Charcot-Marie-Tooth disease type 2Q (CMT2Q, MIM 615025) and eosinophilic esophagitis (EoE) (10–12). There is currently no satisfactory pathophysiological mechanism that explains the observed pleiotropy of *DHTKD1* variants. Structural studies that allow modeling the consequences of the observed amino acid substitutions may help to address this question. Homologous structures have been resolved for *Escherichia coli* E1o (sucA) and *Mycobacterium smegmatis* KGD (2-oxoglurate decarboxylase) (13–15), which respectively share 40% and 37% identity with DHTKD1. Both proteins adopt the common fold of thiamin diphosphate (ThDP)-dependent 2-oxo acid dehydrogenases (16, 17).

The function of OADHc is upstream of glutaryl-CoA dehydrogenase, the enzyme defective in glutaric aciduria type 1 (GA1; MIM 231670) (18–21). The limited clinical consequences of AMOXAD has led multiple investigators to hypothesize that inhibition of DHTKD1 is a potential therapeutic strategy to treat GA1 (2, 8, 22). Unfortunately, the substantial substrate overlap between OGDHc and OADHc precludes a significant reduction in the accumulation of toxic substrates (8, 22). Nevertheless, inhibitors are useful tools that will help shed light on this important metabolic pathway and can help us understand its role in several human diseases. Here we report the crystal structure of the first human DHTKD1 at 2.1 Å in complex with ThDP and identify the first drug-like inhibitors of the enzyme.

## Results

### Identification of substrate analogs as DHTKD1 inhibitors

In order to be able to screen for DHTKD1 inhibitors, we optimized an assay based on dichlorophenolindophenol (DCPIP) reduction (Figure S2a). We first assessed substrate specificity of DHTKD1 by measuring the activity with 2-oxoglutaric acid (OG), OA, 2-oxopimelic acid (OP) and 2-oxosuberic acid (OS), which vary only in carbon chain length (Figure 1). Consistent with the genetic evidence and work by others (2, 4, 7), we found that DHTKD1 has optimal activity with the canonical substrate OA (Figure S1) and negligible activity with OG and OS. Interestingly, DHTKD1 also displayed activity with OP, a metabolite currently unknown in human metabolism. Although the affinity of DHTKD1 for OP was lower than for OA, the V_max_ appeared higher with OP (Figure 1a). As judged from the progress curves, the assay was nonlinear with initial velocity being the highest, which is common for these DCPIP-based assays. Remarkably, the activity with OA declined much more rapidly than the activity with OP (Figure S2b). Therefore, OP provides a better dynamic window in inhibitor screening assays.

**Figure 1.**
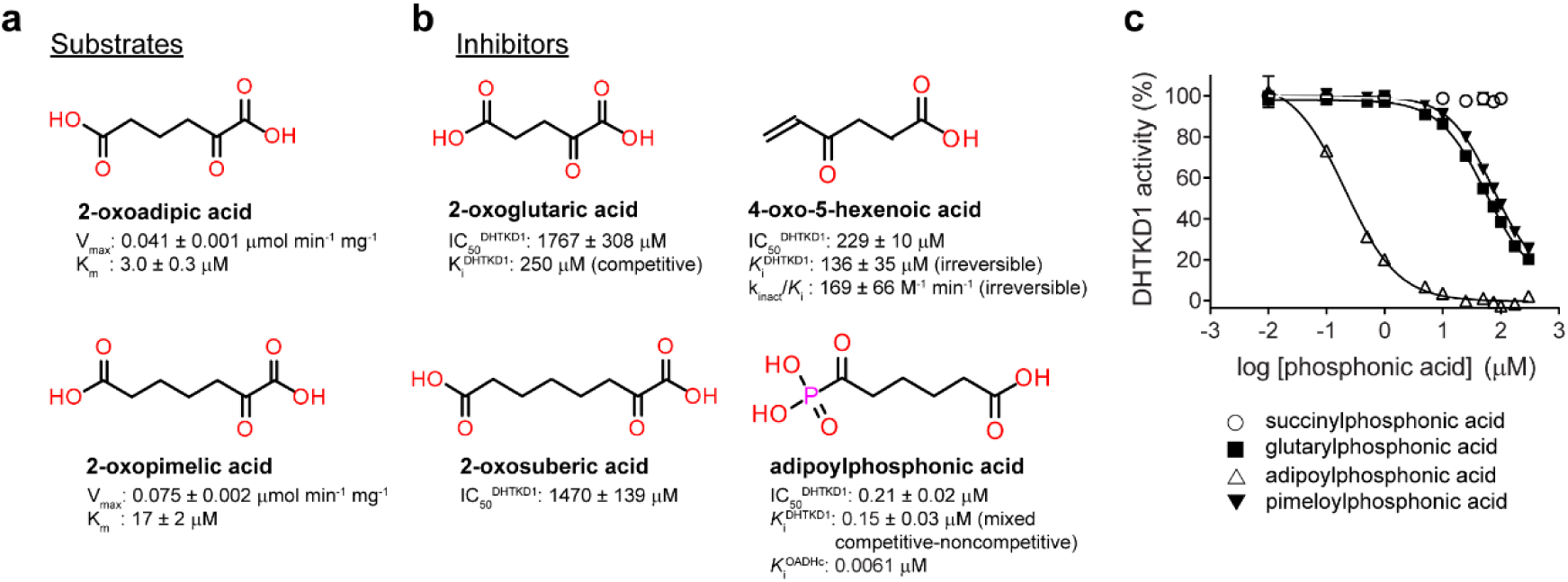
Substrates and inhibitors of DHTKD1. (a) Structure of DHTKD1 substrates and their kinetic parameters. (b) Structure and IC_50_, *K*_i_ and k_inact_/*K*_i_ (for irreversible inhibitors) of DHTKD1 inhibitors. (c) Dose-response curves of DHTKD1 inhibition by phosphonic acids, using OP as substrate.

Six out of the 23 selected substrate analogs were able to inhibit DHTKD1 (Table S1). OG and OS inhibited DHTKD1 activity competitively (Figure 1b, Figure S2c and Table S1). These data indicate that while DHTKD1 prefers OA and OP as substrates, OG and OS are able to bind to the enzyme, which is consistent with previous work (4). Vigabatrin, an anti-epileptic and irreversible transaminase inhibitor known to cause α-aminoadipic aciduria (23), did not inhibit DHTKD1, but its oxo analog 4-oxo-5-hexenoic acid inhibited DHTKD1 irreversibly (Figure 1b). OG, OS and 4-oxo-5-hexenoic acid had relatively low affinities for DHTKD1.

Phosphonic acids inhibit 2-oxo acid dehydrogenase complexes by forming a pre-decarboxylation nucleophilic adduct at the C2 thiazolium atom, the adduct resembling a transition state analog (24–26). We evaluated the ability of succinyl-, glutaryl-, adipoyl- and pimeloyl-phosphonic acid to inhibit DHTKD1. Adipoylphosphonic acid was a highly effective and specific mixed competitive-noncompetitive inhibitor of DHTKD1 (Ki of 0.15±0.03μM), whereas glutaryl- and pimeloylphosphonic acid were less effective and succinyl-phosphonic acid was ineffective (Figures 1b and c and Figures S2d−f). We next assessed the ability of adipoylphosphonic acid to inhibit human OGDHc and OADHc containing OGDH and DHTKD1, respectively (Figures S2g−j). Adipoylphosphonic acid inhibited OADHc activity with an estimated K_i_^app^ of 0.105±0.038μM. Taking into account that the inhibitor and substrate bind in the same active centers, the K_i_ was an estimated 1.6 nM indicating a tight-binding inhibitor (Figure 1b and Figure S2i). These data indicate that adipoylphosphonic acid is a potent inhibitor of OADHc and not of OGDHc (Figures S2i and j). This also establishes that it is the E1 component that is subject to inhibition by adipoylphosphonic acid rather than the E2 or E3 components.

We next used circular dichroism (CD) to monitor the formation of 1’,4’-iminophosphonoadipoyl-ThDP at ∼305nm, a reporter of the tetrahedral predecarboxylation intermediate in ThDP-dependent enzymes (25–27). We showed that the positive CD305 signal depended on the concentration of adipoylphosphonic acid (Figures S2k−m) and displayed saturation of the DHTKD1 active centers by adipoylphosphonic acid at a molar ratio of 1.87 (with much less or no saturation of the OGDH active centers (Figures S2n−r)). These data demonstrate stoichiometric binding of adipoylphosphonic acid in the active centers of DHTKD1, a finding that correlates with the loss of OADHc activity. Thus, adipoylphosphonic acid is a mechanism-based, active-center-directed potent, tight-binding inhibitor of DHTKD1. To test whether this compound could inhibit DHTKD1 in cells, we treated HEK-293 cells and looked at the levels of AAA, the transamination product of the DHTKD1 substrate OA (Figure S1). Adipoylphosphonic acid dose dependently increased the level of AAA in HEK-293 cells, demonstrating the successful inhibition of DHTKD1 in a cellular model (Figures S2s and t).

### Identification of tenatoprazole as a DHTKD1 inhibitor through HTS

In order to identify DHTKD1 inhibitors with drug-like properties, we developed and performed a high-throughput screen (HTS, Supplemental Figures S7-S10). The class of compounds that were consistently identified as DHTKD1 inhibitors in the screen were benzimidazole and imidazopyridine proton-pump inhibitors, with tenatoprazole being most potent (IC_50_, 27 ± 1 μM) (Figure 2a). Analysis of several omeprazole analogs, including sulfones and sulfides highlighted the importance of the sulfoxide group in the DHTKD1 inhibition process (Figure S3). Tenatoprazole displayed kinetics most compatible with a noncompetitive inhibition of DHTKD1 (*K*i = 83 ± 34 μM) (Figure 2a and Figures S4a and b). We then analyzed the drug-protein interaction by differential scanning fluorimetry (DSF). Surprisingly, tenatoprazole destabilized the protein, decreasing the melting temperature by over 3 degrees (250 μM) or 5 degrees (500 μM) (Figure 2b and Figure S4c), an effect absent in the presence of dithiothreitol (DTT, Figure S4d). We compared the other omeprazole analogs in the same assay and found that destabilization tracked with the IC_50_ values of the inhibitors, with tenatoprazole showing the largest destabilization of all the compounds (Figure 2b). Tenatoprazole was unable to inhibit OADHc, likely due to the presence of DTT in the assay mixture. Tenatoprazole increased AAA levels (Figure 2c) without major effects on other amino acids indicating the inhibition of DHTKD1 is rather specific (Figure S4e).

**Figure 2.**
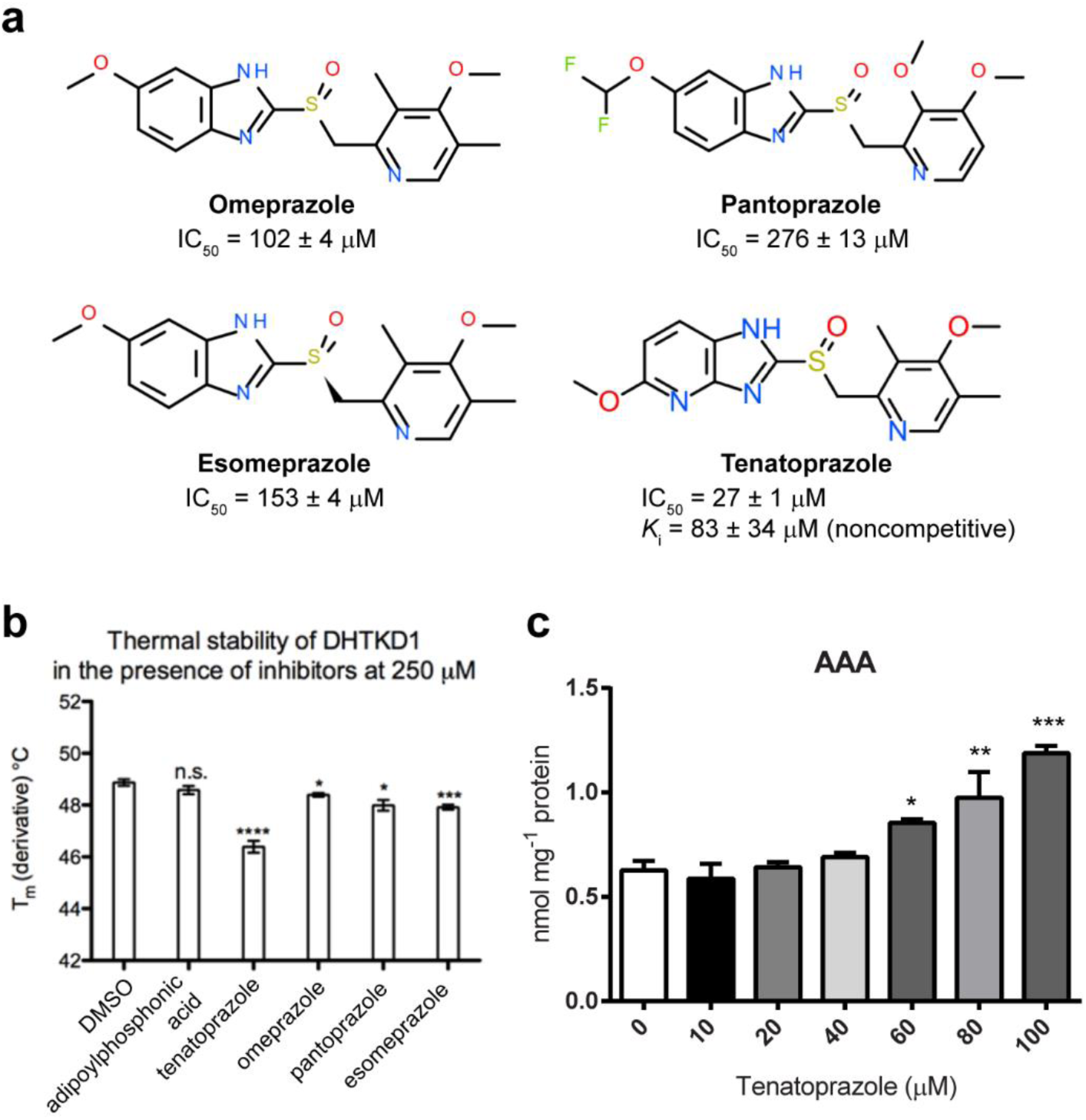
Proton-pump inhibitors are inhibitors of DHTKD1. (a) Structures and IC_50_ of omeprazole, pantoprazole, esomeprazole and tenatoprazole. The K_i_ of tenatoprazole is also shown. (b) Thermal stability of DHTKD1 with proton-pump inhibitors. DHTKD1 protein was incubated with the indicated inhibitors at 250 μM and then analyzed by DSF. Adipoylphosphonic acid was also analyzed. (c) Tenatoprazole inhibits DHTKD1 in HEK-293 cells. HEK-293 cells were incubated with 0-100 μM tenatoprazole and the level of AAA was measured. \**P* < 0.05, \*\**P* < 0.01 and \*\*\**P* < 0.001.

### Structure of DHTKD1

We obtained a crystal structure of human DHTKD1 at 2.1 Å (Figure 3a) using the *M. smegmatis* KGD structure as a search model for molecular replacement. The cofactor ThDP was present and bound to several residues in the active site, making several key presumed hydrogen bonds (Figure 3b). DHTKD1 crystallized as a dimer (Figure 3a and Table S2), similar to the *M. smegmatis* homolog, with the two monomers twisted around each other and two active sites formed at the interface. We believe that the minimal functional unit is a dimer, which is evidenced by several observations. First, we note that each of the two active sites in the dimer comprises residues from both copies of the protein, with one chain providing residues for binding the phosphate moieties and the other for the thiamin (Figure 3b). It is unlikely that a single copy can bind ThDP fully. Secondly, there is a magnesium coordinating E641 from each monomer, which is suggestive of a stable ionic interaction (Figure 3c). Thirdly, we note that the surface area of the interface between the two monomers is extensive. According to the PISA server, the interface covers over 5000 Å^2^ of area with a predicted ΔG of –29.6 kcal/mol, suggestive of a strong interaction between the two copies (28). It is likely that the full complex stabilizes the dimer further.

**Figure 3.**
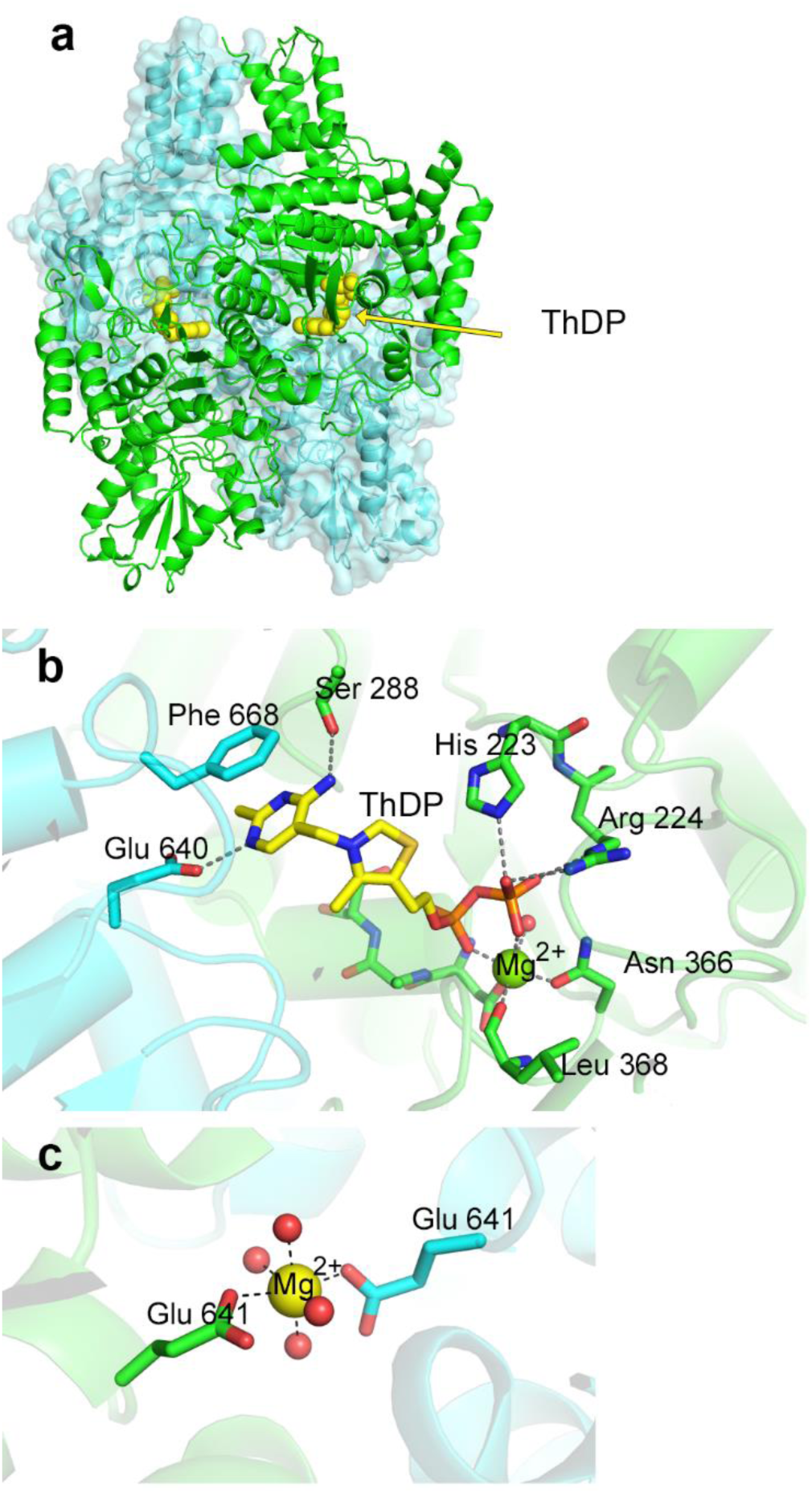
Crystal structure of human DHTKD1 complexed to ThDP. (a) Overall fold of DHTKD1. The dimer is shown with one chain in green and the other in cyan, with the ThDP shown in yellow. (b) Active site of human DHTKD1. Several interactions that help bind the ThDP and magnesium ion are indicated. Residues are colored according to the chain they come from. (c) Dimeric magnesium interface. Coordination of one glutamate from each monomer by a magnesium ion is shown, with the rest of the octahedral magnesium sites coordinated to water.

The tightly packed dimer still leaves room for the substrates to be solvent accessible, so they can be transferred to the other components of the complex (Figure S5a). While the active site is conserved between *M. smegmatis* and human (Figure S5b), we note two major differences. The human protein has a smaller serine (Ser 288) in the active site instead of tyrosine, which provides more room compared to the *M. smegmatis* protein (Figures S5c and d), which may explain the ability of the protein to accept larger substrates such as OA and OP. Additionally human DHTKD1 has a Tyr residue at position 190, which differs from the Phe residue found in the *M. smegmatis* protein and other proteins of the OGDH family, and is predicted to be implicated in substrate binding. We studied the possible binding mode of the preferred DHTKD1 substrates (OA and OP) and the inhibitor adipoylphosphonic acid by docking the compounds into the crystal structure of the protein. Using induced fit docking (IFD), to allow for small adjustments in the protein structure, we obtained a binding mode for OA that is compatible with the formation of a pre-decarboxylation intermediate at the C2 thiazolium atom of ThDP (Figure S5e). The substrate is positioned in the active site by a series of hydrogen bonding interactions with the side chains of Lys 188, Tyr 190, His 435, Asn 436, and His 708. The same interactions were observed in the docking pose of OP (not shown). IFD of the adipoylphosphonic acid inhibitor showed that it mimics the substrate binding mode resulting in a pose that is compatible with the formation of the decarboxylation nucleophilic adduct suggested by the experimental data (Figure S5f).

### Analysis of known DHTKD1 missense variants

Next, we studied all reported missense variants in DHTKD1 (Figure 4a) (7 variants reported in AMOXAD patients (2, 3, 6) and 3 variants associated with EoE (10)) (Figures S6a and b). The S777P and S862I variants were largely insoluble and the small amounts of soluble protein exhibited severe proteolysis during expression and purification. This indicates that the mutations prevent proper folding of the protein. In the crystal structure, the S862 sidechain makes important contacts to sidechains in two different parts of the protein, T917 and H812. Losing these contacts and inserting the bulkier hydrophobic isoleucine likely inhibits protein folding. For S777, there is a likely hydrogen bond between the serine sidechain and the backbone of neighboring L589. The mutation at this position eliminates this interaction as well as introduces a proline, which can destabilize the current backbone conformation due to its different preference in φ,ψ dihedral angles.

**Figure 4.**
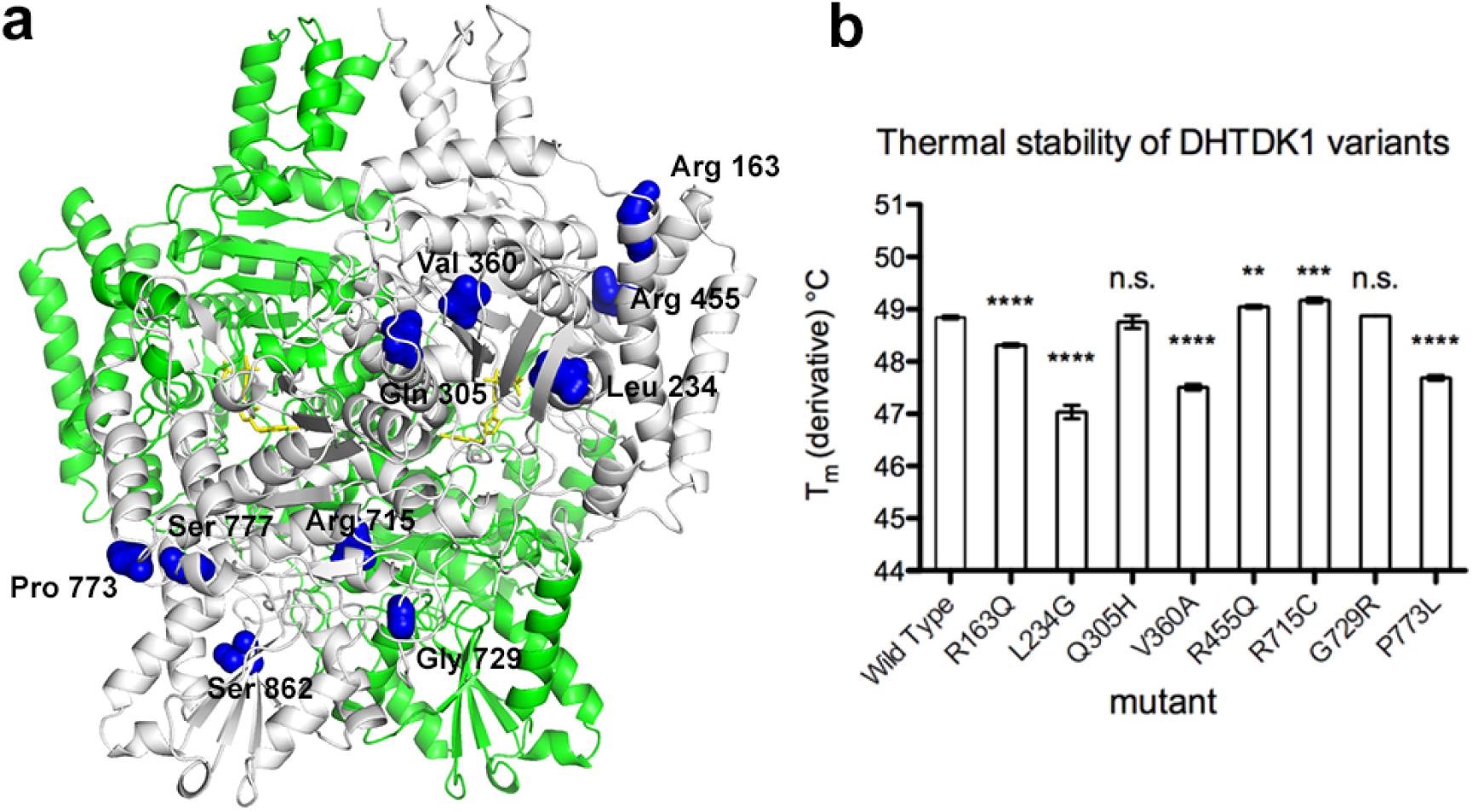
DHTKD1 variants. (a) Structural localization of DHTKD1 missense variants analyzed. The DHTKD1 monomer is shown as a ribbon representation. Amino acid residues affected by mutations are indicated as spheres in dark blue on one monomer. (b) Thermal stability of DHTKD1 mutants analyzed by differential scanning fluorimetry. n=4.

For the 8 other DHTKD1 variants, we analyzed steady-state kinetic parameters and thermal stability (Table 1 and Figure S6c). Three variants displayed a higher propensity to form insoluble protein (L234G, R455Q and R715C), but enough soluble protein was recovered to establish that they were inactive. Two variants (V360A and P773L) showed a decrease catalytic efficiency (1.2- and 2.7-fold decrease, respectively) due to a lower V_max_ when compared with wild-type. Three variants (R163Q, Q305H and G729R) showed kinetic properties similar or slightly improved compared with wild-type (1.3-, 1.7- and 1.6-fold increase in catalytic efficiency, respectively). Substrate affinity was not changed in any of the variants.

**Table 1.** Steady-state kinetic properties of DHTKD1 wild type and variants.

| DHTKD1 | | $V_{\max}$<br>($\mu\text{mol DCPIPH}_2 \cdot \text{min}^{-1} \cdot \text{mg}^{-1}$) | $k_{\text{cat}}$<br>($\text{s}^{-1}$) | $K_m$<br>($\mu\text{M}$) | $k_{\text{cat}}/K_m^a$<br>($\text{M}^{-1} \cdot \text{s}^{-1}$) |
| --- | --- | --- | --- | --- | --- |
| – | WT | $0.029 \pm 0.001$ | 0.097 | $52 \pm 2$ | 1868 |
| c.488G>A | p.R163Q (EoE)<sup>b</sup> | $0.037 \pm 0.001$ | 0.124 | $50 \pm 2$ | 2479 |
| c.700_701delinsGG | p.L234G | *inactive* |  |  |  |
| c.915G>C | p.Q305H | $0.036 \pm 0.001$ | 0.121 | $37 \pm 1$ | 3259 |
| c.1079T>C | p.V360A (EoE) | $0.022 \pm 0.001$ | 0.074 | $46 \pm 3$ | 1602 |
| c.1364G>A | p.R455Q | *inactive* |  |  |  |
| c.2143C>T | p.R715C | *inactive* |  |  |  |
| c.2185G>A | p.G729R | $0.034 \pm 0.001$ | 0.114 | $39 \pm 1$ | 2921 |
| c.2318C>T | p.P773L | $0.011 \pm 0.001$ | 0.037 | $54 \pm 4$ | 682 |
| c.2329 T>C | p.S777P | n.a.<sup>c</sup> |  |  |  |
| c.2585G>T | p.S862I (EoE) | n.a. |  |  |  |

The kinetic properties were determined at 30 °C, 2 mM MgCl<sub>2</sub>, 1 mM ThDP (OP variable). <sup>a</sup>The catalytic efficiency was calculated on the basis of dimer molecular mass of 201 kDa for the DHTKD1 processed form after removal of the mitochondrial transit peptide. $K_m$ represents the OP concentration at half-maximal activity and $k_{\text{cat}}/K_m$ is here defined as the catalytic efficiency. <sup>b</sup>Mutations detected in patients with eosinophilic esophagitis (EoE); all others were detected in 2-aminoadipic and 2-oxoadipic aciduria (AMOXAD) patients. <sup>c</sup>n.a., not available due to the lack of solubility of the full-length protein.

The observed DHTKD1 variants are spread throughout the protein and generally avoid the ThDP binding site (Fig. 4a). All 8 soluble variants were stable enough to determine the melting temperature by DSF testing (Figure 4b). Q305H and G729R had statistically insignificant changes on the Tm of the protein suggesting that their effects are not structural. L234G, V360A, and P773L had decreased T_m_ values. V360 is in the core of the protein and reducing its size likely destabilizes the hydrophobic packing of the protein as observed in other enzymes (29). L234G and P773L likely cause problems in folding by altering to glycine or from proline, which have unique dihedral angle preferences as mentioned above. The remaining 3 variants had relatively small, but reproducible changes with an increase in stability for R455Q and R715C, and a decrease in stability for R163Q. R455 forms a salt bridge with D400 and a hydrogen bond with T430 while R715 makes a salt bridge with D677 and a hydrogen bond with Q673. Mutation of these residues likely abrogates activity by disrupting these important interactions. For Q305H, G729R, but also R163Q, we can likely rule out protein folding as causative. Given that these variants are active, we speculate that they affect assembly or substrate exchange with the E2 or E3 components of OADHc. The G729 and R163 residues are indeed on the surface. Q305 is near the surface but makes important internal contacts, so it is unclear how it could maintain activity, but perhaps the substitution to His maintains these contacts but interferes with the complex.

## Discussion

Here we employed a DHTKD1 specific enzyme assay to enable screening for inhibitors. We found that not only OA, but also OP are good substrates for DHTKD1. The affinity for OA was higher, whereas the V_max_ was higher with OP. Virtually no activity was observed with OG and OS. Interestingly, in vitro reconstituted OADHc (DHTKD1, DLST and DLD) did show activity toward OG (4). This apparent inconsistency is likely explained by differences in the assayed enzyme reactions. OADHc relies on the E2 component DLST for oxidation and transfer of the enamine intermediate to lipoic acid and ultimately CoA. This reaction will not only be determined by the substrate specificity of DHTKD1, but also by that of DLST. Given the canonical role of DLST in the OGDHc, it should work well with succinyl intermediates (3-carboxy-1-hydroxypropyl-ThDP) possibly facilitating DHTKD1 to convert OG. Binding of OG to DHTKD1 is supported by the observation that it acts as a competitive inhibitor in the DHTKD1 specific enzyme assay.

Succinylphosphonic acid is a known inhibitor of OGDHc (4, 30). We have chemically synthesized adipoylphosphonic acid, which is the phosphonic acid analog of the DHTKD1 substrate OP. We have shown that adipoylphosphonic acid acts as a mechanism-based, active site directed tight-binding inhibitor of OADHc, and a much weaker one of the OGDHc. Adipoylphosphonic acid lacks chemical properties to be developed as a drug, but is a molecule that can be used *in vitro* and in whole cell systems. Through HTS, we identified the class of benzimidazole/imidazopyridine proton-pump inhibitors as inhibitors of DHTKD1. Tenatoprazole is a noncompetitive inhibitor of DHTKD1, possibly binding to an allosteric site in the enzyme. We note that tenatoprazole causes an unusually large destabilization of DHTKD1 as measured by DSF, which indicates structural alterations upon binding. We were unable to co-crystallize tenatoprazole with DHTKD1 in the same crystal form consistent with the possibility that tenatoprazole alters the conformation of the protein.

Whereas structures homologous to DHTKD1 have been reported for distant species such as *M. smegmatis* (KGD) and *E. coli* (SucA), we report the first eukaryotic structure for an E1 enzyme. The human structure provides valuable information on the possible mechanism of amino acid substitutions associated with human disease. We expressed all missense mutations that have been associated with AMOXAD and EoE. Five out of the seven DHTKD1 autosomal recessive missense variants associated with AMOXAD showed defects in folding, function or thermal stability. S777P was insoluble, L234G, R455Q and R715C were inactive and P773L presented a lower catalytic activity and thermal stability. Fibroblasts from an AMOXAD patient with compound heterozygosity for R455Q/S777P did not display any immunoreactivity for DHTKD1 (3), which is consistent with our analysis of these two variants. The Q305H and G729R variants, however, did not display any defects in folding, activity and thermal stability. Other possible mechanisms not tested in our studies include a negative impact of the mutation on the assembly with other components of the complex or in the substrate exchange between these components.

The three autosomal dominant missense variants associated with EoE have quite distinct properties. R163Q behaves similar to the wild type protein, V360A had a lower catalytic efficiency and thermal stability and S862I was insoluble. EoE is a chronic allergic inflammatory esophageal disorder and it remains unclear how DHTKD1 variants can produce a dominant effect leading to mitochondrial dysfunction as implicated in the pathogenesis of this disease (10). A possible effect on the assembly of the OADHc and the recently described hybrid complex with OGDHc (8) remains to be demonstrated. Alternatively, the EoE mutations could lead to a conformation-driven gain-of-function mechanism, as has recently been demonstrated for mutations in the aminoacyl-transfer RNA (tRNA) synthetases associated with CMT (31).

In summary, we determine the crystal structure of human DHTKD1, an enzyme associated with AMOXAD, CMT2Q and EoE, which are 3 clinically distinct disorders. We have identified adipoylphosphonic acid and proton-pump inhibitors as specific inhibitors of this protein. These compounds serve as a starting point for more selective and potent inhibitors that can help shed light on the function of this enzyme *in vivo*. Lastly, we have determined the consequences of all DHTDK1 missense variants reported to date, which combined with the human structure, allowed us to begin to understand the role of DHTKD1 in several human diseases.

## Materials and Methods

### Materials

2-oxoheptanedioic acid (2-oxopimelic acid; OP) was obtained from AKos GmbH, 4-oxo-5-hexenoic acid was obtained from Santa Cruz Biotechnology and succinylphosphonic acid was from MedChem Express. DCPIP, ThDP, coenzyme A (CoA), NAD, β-mercaptoethanol and Prionex® were from Millipore-Sigma. HEK-293 cells were obtained from American Type Culture Collection (Manassas, VA, USA). DMEM, penicillin, streptomycin and fetal bovine serum were obtained from Thermo Fisher Scientific (Waltham, MA, USA). All other chemicals were of analytical grade.

### Phosphonic acid analogs synthesis

Reactions in anhydrous solvents were carried out in glassware that was flame-dried or oven-dried. Unless noted, reactions were magnetically stirred and conducted under an atmosphere. Air-sensitive reagents and solutions were transferred via syringe and were introduced to the reaction vessel through rubber septa. Solids were introduced under a positive pressure of Ar. Temperatures, other than room temperature (rt); refer to bath temperatures unless otherwise indicated. All commercially obtained solvents and reagents were used as received. Deionized water was used for all aqueous reactions, work-ups, and for the preparation of all aqueous solutions. The phrase “concentrated in vacuo” refers to removal of solvents by means of a Büchi rotary-evaporator attached to a variable vacuum pump followed by pumping to a constant weight (< 1 Torr). Proton and carbon nuclear magnetic resonance (NMR) spectra were obtained on a Bruker Avance 600 (600 MHz). Chemical shifts are reported in ppm (δ). ^1^H NMR data are reported as follows: chemical shift (multiplicity, coupling constant (Hz), number of hydrogens). Multiplicities are denoted accordingly: s (singlet), d (doublet), dd (doublet of doublets), ddd (doublet of doublet of doublets), dt (doublet of triplets), tt (triplet of triplets), dq (doublet of quartets), t (triplet), q (quartet), p (pentet), m (multiplet). High resolution mass spectra (LCMS) were obtained using an Agilent 1200 Series Rapid Resolution LC/MS. The chromatography was performed by using Teledyne ISCO RediSep normal phase (40-60 microns) silica Gel disposable flash columns using a Teledyne ISCO Combiflash Rf purification system (for detailed synthesis and NMR spectra see supporting information).

### Cloning of DHTKD1 into the pET28a(+) vector

Human DHTKD1 cDNA clone (Genbank accession BC007955) was provided by TransOMIC technologies (Huntsville, AL). The DHTKD1 open reading frame (amino acids 25-919) was amplified using PrimeSTAR GXL DNA Polymerase (Takara) and primers forward (5’-GGT TTA <u>GAA TTC</u> ATG CAG ACC GAG CGG GGC GTT TA-3’) and reverse (5’-CTT TAC <u>CTC GAG</u> TCA TCA AGC GAA GGT CTT GGC GAG GAT-3’). These primers introduce an EcoRI and XhoI site (underlined). The pET-28a(+) vector (Novagen, EMD) was digested with EcoRI-HF and XhoI (NEB), treated by Calf Intestinal Alkaline Phosphatase (NEB) and purified by PureLink Quick Gel Extraction and PCR Purification combo kit (Invitrogen). The DHTKD1 PCR product was also digested with EcoRI-HF and XhoI, purified and ligated into the digested pET-28a(+) vector by T4 DNA Ligase (NEB). DH5α competent cells were transformed with the ligated product and plasmid DNA from single clone was purified and sequence verified. The resulting pET-28a(+)-DHTKD1 plasmid encodes an N-terminal His-tagged human DHTDK1 suitable for protein production.

### Protein expression and purification of wild type DHTKD1

Rosetta 2(DE3)pLysS competent cells (Novagen, EMD) were transformed with the pET-28a(+)-encoding wild type DHTKD1 (amino acids 25-919) and single clones were used for protein production. Cells were grown in LB medium with kanamycin (25 μg/ml for pET-28a) and chloramphenicol (35 μg/ml for Rosetta cells). A culture for protein production was inoculated 1 in 1000 with a fresh overnight culture and cells were grown at 37°C with shaking for 3-4 h with regular determination of the optical density at 600 nm (OD600). Once the culture reached an OD600 of 0.6-0.7, the culture was placed at RT, and protein production was induced by adding IPTG to a final concentration of 1 mM. Protein production was continued overnight, after which the cells were harvested by centrifugation and stored at -20 °C. Protein extracts were prepared using BugBuster (Millipore Sigma) diluted in binding buffer (20 mM Tris-HCl pH 7.8, 500 mM NaCl, 5 mM imidazole, 10 mM β-mercaptoethanol, 50% BugBuster) with 1x cOmplete, mini, EDTA-free protease inhibitor cocktail. The extract was sonicated and the protein purified by IMAC and dialyzed against 100 mM potassium phosphate pH 7, 2.1 mM MgCl_2_, 0.21 mM ThDP, 10% glycerol. The protein was stored at -20 °C in dialysis buffer with 20% glycerol. Protein purity was analyzed by SDS-PAGE in a 4-12% (w/v) polyacrylamide gel. Protein concentration was estimated by the Bradford method (32) using bovine serum albumin (BSA) as standard.

### DHTKD1 specific enzyme activity assay

In order to measure activity of the isolated DHTKD1 E1 component and to be able to screen for DHTKD1 inhibitors, we have adapted an assay using the redox dye 2,6-dichlorophenolindophenol (DCPIP), which oxidizes the enamine thiamin diphosphate (ThDP) intermediate and has been used for many years in the ThDP field (33, 34). The assay was optimized for pH, KCl, ThDP, MgCl_2_, detergent and carrier protein concentration. The established assay conditions are 50 mM MOPS pH 7.4, 100 mM KCl, 2 mM MgCl_2_, 1 mM ThDP, 0.1 mM DCPIP, 0.1% Triton X-100 and 0.25% Prionex. Recombinant DHTKD1 E1 component was used at 0.025 mg/mL and OP was used at 62.5 μM as substrate, unless otherwise indicated.

The kinetics of the reaction was monitored as the decrease in absorption at 600 nm using 96-well plates and an EnVision Multilabel Plate Reader (PerkinElmer) or a similar plate reader. A substrate blank for correction of DCPIP reduction was included in all assays and subtracted. The millimolar extinction coefficient used for DCPIP was 20.6 mM^-1^ cm^-1^ (35). The stock solutions containing the substrates OG, OA, OP and OS were neutralized using KOH. Compounds were dissolved in DMSO and tested at 300 μM (1% DMSO final concentration), unless otherwise stated. IC_50_ was determined for selected compounds using the GraphPad Prism 7 software. Steady-state kinetic data were analyzed by nonlinear regression analysis using GraphPad Prism 7 software and a mixed model inhibition. The irreversible inhibitor 4-oxo-5-hexenoic acid was evaluated by the second-order rate constant k_inact_/*K*_i_ that describes the overall potency of inactivation, where *K*_i_ is the inhibition constant of the first reversible binding event and k_inact_ is the maximum rate of inactivation (36). Progress curves were fitted to a single exponential equation to obtain the observed first order rate constant of inactivation, k_obs_. The dependence of k_obs_ on inhibitor concentration was used to calculate the k_inact_/*K*_i_ parameter.

### OGDHc activity

OGDHc from porcine heart (Sigma K1502) was mixed with assay buffer (final concentration: 50 mM MOPS, pH 7.4, 0.2 mM MgCl_2_, 0.01 mM CaCl_2_, 0.3 mM ThDP, 0.12 mM CoA, 2 mM NAD, 2.6 mM β-mercaptoethanol). The reaction was started by the addition of 1 mM substrate OG. The activity of OGDHc was followed by measuring the NADH production at 340 nm at 30 °C and steady-state velocities were taken from the linear portion of the time curve.

### Overall OADHc activity of NADH production *in vitro*

The OADHc was assembled from recombinant DHTKD1, DLST (E2) and DLD (E3) components expressed and purified independently. For OADHc assembly, the DHTKD1 (0.4 mg) in 0.1 M Tris.HCl (pH 7.5) containing 0.3 M NH4Cl, 0.5 mM ThDP and 2.0 mM MgCl_2_ was mixed with DLST (0.80 mg) and DLD (2.0 mg) at a mass ratio (mg/mg/mg) of 1:2:5 and was incubated 40 min at 25 °C. An aliquot containing 0.01 mg of DHTKD1 and the corresponding amounts of E2 and E3 was withdrawn and was placed into 1 ml of the reaction assay containing all components necessary for the overall OADHc activity assay (4, 5, 37). The reaction was initiated by addition of OA (1.0 mM) and CoA (400 μM) and the progress curves were recorded at 37 °C for 3 min. The reaction rates (slope/min at 340 nm) were calculated from the linear part of the recorded progress curves.

### Circular dichroism analysis of 1’,4’-iminophosphonoadipoyl-ThDP intermediate

CD spectra of DHTKD1 (19.4 μM concentration of active centers) in 100 mM HEPES (pH 7.5) containing 0.5 mM ThDP, 2.0 mM MgCl_2_, 0.15 M NaCl and 5% glycerol were recorded in the absence and in the presence of adipoylphosphonic acid (1-51 μM) on a Chirascan CD spectrometer (Applied Photophysics, Leatherhead, UK) in 1-cm path length cell in the near-UV CD region (290-450 nm) at 25 °C. The intensity of the CD band at 305 nm was estimated and was plotted versus the concentration of adipoylphosphonic acid or *versus* the [adipoylphosphonic acid]/[DHTKD1 active centers] ratio.

### High-throughput screening

In order to identify DHTKD1 inhibitors with drug-like properties, we developed and performed a high-throughput screen based on the DHTKD1 activity assay (Figures S2a and S7a). We evaluated the robustness of the assay by performing a pilot screen of ∼4000 molecules from a drug and bioactive collection and ∼600 molecules from a diversity collection (ChemBridge). The screen was robust with a good assay window and percent coefficient of variance % CV < 5% and Z′ of 0.84 ± 0.10 (Figure S7b) (38). When considering compounds with inhibition greater than 3SD above the mean, the hit rate for drug (38 hit compounds, 1.7%) and bio-actives (26 hit compounds, 1.4%) was significantly higher when compared to the lower hit rate for diversity molecules (5 hit compounds, 0.8%) (Figures S7c and d).

We then screened against a diversity library, a collection of 125,465 compounds from three libraries (ChemBridge, ChemDiv and CB Premium). Using a threshold criterion for inhibition greater than or equal to 3SD above the mean (Bscore, inhibition and standard score), we identified 320 hit compounds (pilot screen and HTS) (Figure S8a). This group included 257 singletons and 63 compounds in 26 clusters (≥2 members with similar structure) and corresponds to a hit rate of ∼0.25%. Additional filtering resulted in a final number of 133 hit compounds (106 singletons and 27 compounds in 11 clusters, a hit rate of ∼0.17%) (Figure S8b). We then selected 192 hit molecules from the primary screen (including all 133 hits after row exclusion and 59 after additional review but before row exclusion) for testing in a concentration-response inhibition assay. 40 out of the 192 hit molecules showed dose-response curves (Figure S9). 16 of these molecules had IC_50_ ≤30 μM, and 13 were commercially available and advanced to the repurchase and re-evaluation phase (Figure S10).

The HTS was conducted by transferring fifty nanoliters of compound in DMSO (10 μM final compound concentration, 0.1% DMSO) by pin tool to 30 µL of the reaction mixture A (buffer assay mixture and substrate) with shaking (Beckman Biomek FXP, Brea, CA). The activity assay was started by adding reaction mixture B (enzyme in storage buffer; 20 μL). The reaction proceeded at room temperature for 30 min and was quenched by adding 100 μM of ZnCl2, an inhibitor of DHTKD1 (IC_50_ = 2.7 ± 0.1 μM). The 384-well plates were read on an EnVision plate reader (PerkinElmer, Waltham, MA; absorbance at 600 nm). Each plate included mock (no enzyme) and positive (adipoylphosphonic acid) controls. Assay performance was measured by z-prime (Z′), a dimensionless factor used to assess the quality of a HTS (38). Hit selection was based on dependent (% inhibition) and independent (B score and standard score) activity parameters. The hit selection threshold was set at calculate mean + 3SD and candidates must satisfy statistical criteria for all three parameters. Standard score (or z-score: (inhibition-mean)/SD)) and Bscore were measured as defined by Brideau et al. (39). Additional further filtering of the screening hits was done by removing systematic influences (e.g. pattern of high absorbance/apparently inhibited rows). Compounds identified in the primary screen were cherry picked for dose-response assays. 150 nL of each compound was transferred by pin tool to each well with concentration ranging from 58 nM to 30 μM. Primary screen results were analyzed using the UCSF SMDC web-based application HiTS. This web-based analysis platform provides a data analysis interface for viewing primary screen and dose-response results. Data analysis for dose-response inhibition experiments was carried out in GraphPad Prism 7 and curves were fit to a standard inhibition log dose-response curve to yield an IC_50_ value.

### Assessment of DHTKD1 inhibitors in HEK-293 cells by amino acid analysis

HEK-293 cells (ATCC CRL-1573) were cultured in DMEM with 4.5 g/L glucose, 584 mg/L L-glutamine and 110 mg/L sodium pyruvate, supplemented with 10% FBS, 100 U/mL penicillin, 100 μg/mL streptomycin, in a humidified atmosphere of 5% CO_2_ at 37 °C. For quantification of 2-aminoadipic acid (AAA) in HEK-293, the cells were incubated in DMEM supplemented with inhibitors using 0.1% DMSO or DMEM as the vehicle control for 24 h. Cells were collected, washed in PBS, flash-frozen in liquid nitrogen and stored at -80°C until further use. Samples were sonicated in ice cold demineralized water and then immediately deproteinized by adding ice cold acetonitrile up to 80% (v/v). After 10 min centrifugation at maximum speed (4 °C), the supernatant was dried under nitrogen and the residue was dissolved in 75 μL of demineralized water. Amino acids were measured by LC-MS/MS (Sema4 formerly the Mount Sinai Genetic Testing Lab) as described (40). Protein pellets were dissolved in 25 mM KOH containing 0.1% Triton X-100, and the protein amount determined by the BCA method for subsequent normalization of the amino acid levels.

### Protein purification for crystallography

*DHTKD1* was cloned into a pET24 vector with a C-terminal 7x His tag. The plasmid was transformed into *LOBSTR*-BL21(DE3), a low background strain of *E. coli*. An overnight culture was used to inoculate 1 L of TB medium (1:1000) containing 50 μg/ml of kanamycin and grown at 37 °C until the cell density reached an OD600 of 1.0. The cells were then cooled to 16 °C and induced overnight with 0.4 mM isopropyl-ß-D-thiogalactopyranoside (IPTG). The next day, cells were harvested and resuspended in tris-buffered saline (20 mM Tris pH 8.0, 250 mM NaCl) and lysed with a french pressure cell at 25K–30K p.s.i. After pelleting down the cellular debris, the lysate was loaded onto a column containing HisPur^TM^ Ni-NTA Resin (Thermo scientific) for immobilized metal affinity chromatography (IMAC) purification. The column was equilibrated with 7 column volumes of TBS supplemented with 40 mM imidazole pH 8.0 then washed with 7 column volumes of 50 mM Hepes buffer pH=7.5 – 7.6 containing 150 mM NaCl, 25 mM imidazole and adjusted to pH 7.5. The column was then eluted step-wise with an elution buffer containing 20 mM Tris-HCl pH = 8.0, 2 mM MgCl_2_, 2 mM ThDP and 250 mM imidazole pH = 8.0 in 10% glycerol. The elution fractions were then pooled after the presence of protein was confirmed using SDS-PAGE. The pooled fractions were then dialyzed against a dialysis buffer (50 mM Hepes pH = 7.6, 150 mM NaCl, 2mM MgCl_2_, 1 mM ThDP) at 4 °C using a 20 kDa MWCO dialysis cassette (Thermo Scientific).

The protein did not produce diffracting crystals initially, so we then methylated the protein to improve crystal packing. Lysines were methylated following the method of Walter et al (41). 20 μL of freshly prepared 1 M dimethylamine-borane complex (DMAB) and 40 μL of 1 M formaldehyde (HCHO) were added to 1 ml of 10 mg/ml of DHTKD1 in a 1.5 ml Eppendorf tube. The tube was covered with aluminum foil and the reactions were gently mixed and incubated for 2 hrs in cold room in that covered tube. Addition of DMAB and HCHO followed by 2 hrs of incubation were repeated twice. Finally, 10 μL of DMAB was added and the reaction was incubated overnight at 4 °C in the cold room. The next day, the methylation reaction was quenched by adding 125 μL of 1 M Tris, pH 7.5. Occasionally, this reaction caused some amount of proteins to be precipitated, which was removed by high speed centrifugation prior to size exclusion chromatography. Following methylation, the protein was then purified by gel filtration on a Superdex 200 increase column (GE Lifescience) in a buffer of 20 mM Tris-HCl pH 8.0, 100 mM KCl, 1 mM MgCl_2_, 1 mM DTT and 0.5 mM ThDP. Protein was then concentrated to approximately 10 mg/ml and flash frozen in liquid nitrogen.

### Crystallization and structure determination

After screening crystal conditions, crystals were grown by the hanging drop vapor diffusion method by mixing 1 μl protein with 1 μl of reservoir containing 0.1 M magnesium acetate, 0.1 M MOPS pH 7.5, and 12% PEG 8000 and 10% glycerol. Crystals were flash frozen in liquid nitrogen before 24 collecting at the AMX (17-ID-1) beamline at NSLS II at Brookhaven National Laboratory. Data was processed using autoproc (42) and scaled with Aimless (43). The structure was solved by molecular replacement, using the structure of the *M. smegmatus* 2-oxoglutarate dehydrogenase protein as a search model (PDB code 3ZHQ) using the program Phaser (44) after making a homology model using Phenix Sculptor (45). After molecular replacement, we performed multiples rounds of building using Phenix Autobuild (46). The models were subsequently refined using Phenix (47), with rigid body refinement and multiple rounds of simulated annealing, minimization, atomic displacement parameter (ADP or B-factor) refinement and TLS refinement (determined using the TLSMD server (48, 49)), with interspersed manual adjustments using Coot (50). Geometric restraints for N-methyl-lysine were generated with Phenix Elbow (51), and these restraints were used throughout refinement. All structural figures were made with Pymol (52).

### Ligand Docking

The crystal structure of DHTKD1 was prepared for docking using the Protein Preparation wizard in Maestro (Schrödinger) to add missing atoms, optimize hydrogen bonds, and minimize the structure (heavy atom convergence to RMSD 0.3 Å). Structures of ligands were prepared using ligprep (Schrödinger) with the OPLS3e force field and Epik ionization. Induced Fit Docking was carried out with Glide/Prime (Schrödinger) using the standard protocol and docking box centered on the active site of the crystal structure. Figures were generated using PyMOL (Schrödinger).

### Site-specific mutagenesis and protein expression and purification of DHTKD1 variants

The DHTKD1 variants were constructed using the QuikChange^®^ II site-directed mutagenesis kit (Thermo Fisher Scientific Inc., MA, USA), the pET28a(+) vector encoding DHTKD1 as template and specific oligonucleotide primers listed in Table S3. Authenticity of the mutagenesis was verified by DNA sequencing.

### Differential scanning fluorimetry

Differential scanning fluorimetry (DSF or thermal shift assay) for the DHTKD1 and its variants was performed on QuantStudio 3 real-time PCR system (Applied Biosystems). Purified proteins were diluted to a final concentration of 0.5 mg/ml in 100 mM potassium phosphate, pH 7, 2.1 mM MgCl_2_, 0.21 mM ThDP, 20% glycerol. 17.5 μL of this 0.5 mg/ml protein was mixed with 2.5 μL of 8X SYPRO orange (Applied Biosystems) in a final 20 μL reaction volume. This 8X SYPRO orange was prepared by diluting the manufacturer supplied dye 250-fold in H_2_O. The reactions were kept at 25 °C for 2 min and then heated from 25 to 95 °C with a rate of 0.05 °C/s. The change in the fluorescence intensities of SYPRO orange was monitored as a function of the temperature and analyzed by Protein Thermal Shift Software 1.3. Each reaction was performed in 5 replicates. For substrates of DHTKD1 it was identical, except there were 4 replicates.

### Statistics

Data are displayed as the mean ± the standard deviation (SD). Differences between groups were evaluated using a one-way analysis of variance with Bonferroni’s multiple comparison test (GraphPad Prism 7). Significance is indicated as follows: \**P* < 0.05, \*\**P* < 0.01, \*\*\**P* < 0.001 and \*\*\*\**P* < 0.0001.

## Accession codes

The structure of DHTKD1 bound to ThDP and Mg^2+^ have been deposited with the Protein Data Bank as 6U3J.

## Acknowledgments

Research reported in this publication was supported by the Eunice Kennedy Shriver National Institute of Child Health & Human Development of the National Institutes of Health under Award Number R03HD092878 (to S.M.H.) and R21HD088775 (to S.M.H. and R.J.D.) and R35GM124838 (to M.B.L.) and by grant UL1TR001433 from the National Center for Advancing Translational Sciences, National Institutes of Health.

This research used resources of the National Synchrotron Light Source II, a U.S. Department of Energy (DOE) Office of Science User Facility operated for the DOE Office of Science by Brookhaven National Laboratory under Contract No. DE-SC0012704. The Life Science Biomedical Technology Research resource is primarily supported by the National Institute of Health, National Institute of General Medical Sciences (NIGMS) through a Biomedical Technology Research Resource P41 grant (P41GM111244), and by the DOE Office of Biological and Environmental Research (KP1605010).

## References

1. V. I. Bunik, D. Degtyarev, Structure-function relationships in the 2-oxo acid dehydrogenase family: substrate-specific signatures and functional predictions for the 2-oxoglutarate dehydrogenase-like proteins. Proteins 71, 874–890 (2008).

2. K. Danhauser et al., DHTKD1 mutations cause 2-aminoadipic and 2-oxoadipic aciduria. Am. J. Hum. Genet. 91, 1082–1087 (2012).

3. J. Hagen et al., Genetic basis of alpha-aminoadipic and alpha-ketoadipic aciduria. J. Inherit. Metab. Dis. 38, 873–879 (2015).

4. N. S. Nemeria et al., The mitochondrial 2-oxoadipate and 2-oxoglutarate dehydrogenase complexes share their E2 and E3 components for their function and both generate reactive oxygen species. Free Radic. Biol. Med. 115, 136–145 (2018).

5. N. S. Nemeria, G. Gerfen, L. Yang, X. Zhang, F. Jordan, Evidence for functional and regulatory cross-talk between the tricarboxylic acid cycle 2-oxoglutarate dehydrogenase complex and 2-oxoadipate dehydrogenase on the l-lysine, l-hydroxylysine and l-tryptophan degradation pathways from studies in vitro. Biochim. Biophys. Acta Bioenerg. 1859, 932–939 (2018).

6. A. R. Stiles et al., New Cases of DHTKD1 Mutations in Patients with 2-Ketoadipic Aciduria. JIMD Rep. 25, 15–19 (2016).

7. Y. Wu et al., Multilayered genetic and omics dissection of mitochondrial activity in a mouse reference population. Cell 158, 1415–1430 (2014).

8. J. Leandro et al., DHTKD1 and OGDH display in vivo substrate overlap and form a hybrid ketoacid dehydrogenase complex. bioRxiv https://doi.org/10.1101/645689 (2019).

9. S. I. Goodman, M. Duran, “Biochemical phenotypes of questionable clinical significance” in Physician’s guide to the diagnosis, treatment, and follow-up of inherited metabolic diseases, N. Blau, M. Duran, K. M. Gibson, C. Dionisi-Vici, Eds. (Springer Verlag: Heidelberg, 2014), pp. 691–705.

10. J. D. Sherrill et al., Whole-exome sequencing uncovers oxidoreductases DHTKD1 and OGDHL as linkers between mitochondrial dysfunction and eosinophilic esophagitis. JCI Insight 3 (2018).

11. W. Y. Xu et al., A nonsense mutation in DHTKD1 causes Charcot-Marie-Tooth disease type 2 in a large Chinese pedigree. Am. J. Hum. Genet. 91, 1088–1094 (2012).

12. M. F. Dohrn et al., Frequent genes in rare diseases: panel-based next generation sequencing to disclose causal mutations in hereditary neuropathies. J. Neurochem. 143, 507–522 (2017).

13. R. A. Frank, A. J. Price, F. D. Northrop, R. N. Perham, B. F. Luisi, Crystal structure of the E1 component of the Escherichia coli 2-oxoglutarate dehydrogenase multienzyme complex. J. Mol. Biol. 368, 639–651 (2007).

14. T. Wagner, N. Barilone, P. M. Alzari, M. Bellinzoni, A dual conformation of the post-decarboxylation intermediate is associated with distinct enzyme states in mycobacterial KGD (alpha-ketoglutarate decarboxylase). Biochem. J. 457, 425–434 (2014).

15. T. Wagner, M. Bellinzoni, A. Wehenkel, H. M. O’Hare, P. M. Alzari, Functional plasticity and allosteric regulation of alpha-ketoglutarate decarboxylase in central mycobacterial metabolism. Chem. Biol. 18, 1011–1020 (2011).

16. A. Ævarsson et al., Crystal structure of human branched-chain alpha-ketoacid dehydrogenase and the molecular basis of multienzyme complex deficiency in maple syrup urine disease. Structure 8, 277–291 (2000).

17. P. Arjunan et al., Structure of the pyruvate dehydrogenase multienzyme complex E1 component from Escherichia coli at 1.85 A resolution. Biochemistry 41, 5213–5221 (2002).

18. N. Boy et al., Patterns, evolution, and severity of striatal injury in insidious-vs acute-onset glutaric aciduria type 1. J. Inherit. Metab. Dis. 42, 117–127 (2019).

19. S. I. Goodman et al., Cloning of glutaryl-CoA dehydrogenase cDNA, and expression of wild type and mutant enzymes in Escherichia coli. Hum. Mol. Genet. 4, 1493–1498 (1995).

20. S. I. Goodman et al., Glutaryl-CoA dehydrogenase mutations in glutaric acidemia (type I): review and report of thirty novel mutations. Hum. Mutat. 12, 141–144 (1998).

21. C. R. Greenberg et al., A G-to-T transversion at the +5 position of intron 1 in the glutaryl CoA dehydrogenase gene is associated with the Island Lake variant of glutaric acidemia type I. Hum. Mol. Genet. 4, 493–495 (1995).

22. C. Biagosch et al., Elevated glutaric acid levels in Dhtkd1-/Gcdh-double knockout mice challenge our current understanding of lysine metabolism. Biochim. Biophys. Acta Mol. Basis Dis. 1863, 2220–2228 (2017).

23. C. Vallat et al., Treatment with vigabatrin may mimic alpha-aminoadipic aciduria. Epilepsia 37, 803–805 (1996).

24. A. V. Artiukhov, A. V. Graf, V. I. Bunik, Directed Regulation of Multienzyme Complexes of 2-Oxo Acid Dehydrogenases Using Phosphonate and Phosphinate Analogs of 2-Oxo Acids. Biochemistry (Mosc.) 81, 1498–1521 (2016).

25. N. Nemeria et al., The 1’,4’-iminopyrimidine tautomer of thiamin diphosphate is poised for catalysis in asymmetric active centers on enzymes. Proc. Natl. Acad. Sci. U. S. A. 104, 78–82 (2007).

26. N. S. Nemeria, S. Chakraborty, A. Balakrishnan, F. Jordan, Reaction mechanisms of thiamin diphosphate enzymes: defining states of ionization and tautomerization of the cofactor at individual steps. FEBS J. 276, 2432–2446 (2009).

27. A. T. Baykal, L. Kakalis, F. Jordan, Electronic and nuclear magnetic resonance spectroscopic features of the 1’,4’-iminopyrimidine tautomeric form of thiamin diphosphate, a novel intermediate on enzymes requiring this coenzyme. Biochemistry 45, 7522–7528 (2006).

28. E. Krissinel, K. Henrick, Inference of macromolecular assemblies from crystalline state. J. Mol. Biol. 372, 774–797 (2007).

29. J. Xu, W. A. Baase, E. Baldwin, B. W. Matthews, The response of T4 lysozyme to large-to-small substitutions within the core and its relation to the hydrophobic effect. Protein Sci. 7, 158–177 (1998).

30. V. I. Bunik et al., Phosphonate analogues of alpha-ketoglutarate inhibit the activity of the alpha-ketoglutarate dehydrogenase complex isolated from brain and in cultured cells. Biochemistry 44, 10552–10561 (2005).

31. D. Blocquel et al., CMT disease severity correlates with mutation-induced open conformation of histidyl-tRNA synthetase, not aminoacylation loss, in patient cells. Proc. Natl. Acad. Sci. U. S. A. 116, 19440–19448 (2019).

32. M. M. Bradford, A rapid and sensitive method for the quantitation of microgram quantities of protein utilizing the principle of protein-dye binding. Anal. Biochem. 72, 248–254 (1976).

33. G. Hubner, A. Schellenberger, R. Bernhardt, S. Khailova, S. E. Severin, Inactivation of the pyruvate dehydrogenase component from pigeon breast muscle pyruvate dehydrogenase complex by alpha-ketobutyric acid. FEBS Lett. 84, 179–182 (1977).

34. L. S. Khailova, R. Bernkhardt, G. Khiubner, [Study of the kinetic mechanism of the pyruvate-2,6-dichlorophenolindophenol reductase activity of muscle pyruvate dehydrogenase]. Biokhimiia 42, 113–117 (1977).

35. J. M. Armstrong, The Molar Extinction Coefficient of 2,6-Dichlorophenol Indophenol. Biochim. Biophys. Acta 86, 194–197 (1964).

36. R. A. Copeland, Evaluation of enzyme inhibitors in drug discovery : a guide for medicinal chemists and pharmacologists (Wiley, Hoboken, N.J., ed. 2nd, 2013), pp. 538 p.

37. N. S. Nemeria et al., The human Krebs cycle 2-oxoglutarate dehydrogenase complex creates an additional source of superoxide/hydrogen peroxide from 2-oxoadipate as alternative substrate. Free Radic. Biol. Med. 108, 644–654 (2017).

38. J. H. Zhang, T. D. Chung, K. R. Oldenburg, A Simple Statistical Parameter for Use in Evaluation and Validation of High Throughput Screening Assays. J. Biomol. Screen. 4, 67–73 (1999).

39. C. Brideau, B. Gunter, B. Pikounis, A. Liaw, Improved statistical methods for hit selection in high-throughput screening. J. Biomol. Screen. 8, 634–647 (2003).

40. A. Le, A. Ng, T. Kwan, K. Cusmano-Ozog, T. M. Cowan, A rapid, sensitive method for quantitative analysis of underivatized amino acids by liquid chromatography-tandem mass spectrometry (LC-MS/MS). J. Chromatogr. B Analyt. Technol. Biomed. Life Sci. 944, 166–174 (2014).

41. T. S. Walter et al., Lysine methylation as a routine rescue strategy for protein crystallization. Structure 14, 1617–1622 (2006).

42. C. Vonrhein et al., Data processing and analysis with the autoPROC toolbox. Acta Crystallogr. D Biol. Crystallogr. 67, 293–302 (2011).

43. P. R. Evans, G. N. Murshudov, How good are my data and what is the resolution? Acta Crystallogr. D Biol. Crystallogr. 69, 1204–1214 (2013).

44. A. J. McCoy et al., Phaser crystallographic software. J. Appl. Crystallogr. 40, 658–674 (2007).

45. G. Bunkoczi, R. J. Read, Improvement of molecular-replacement models with Sculptor. Acta Crystallogr. D Biol. Crystallogr. 67, 303–312 (2011).

46. T. C. Terwilliger et al., Iterative model building, structure refinement and density modification with the PHENIX AutoBuild wizard. Acta Crystallogr. D Biol. Crystallogr. 64, 61–69 (2008).

47. P. D. Adams et al., PHENIX: a comprehensive Python-based system for macromolecular structure solution. Acta Crystallogr. D Biol. Crystallogr. 66, 213–221 (2010).

48. J. Painter, E. A. Merritt, Optimal description of a protein structure in terms of multiple groups undergoing TLS motion. Acta Crystallogr. D Biol. Crystallogr. 62, 439–450 (2006).

49. J. Painter, E. A. Merritt, TLSMD web server for the generation of multi-group TLS models. J. Appl. Crystallogr. 39, 109–111 (2006).

50. P. Emsley, B. Lohkamp, W. G. Scott, K. Cowtan, Features and development of Coot. Acta Crystallogr. D Biol. Crystallogr. 66, 486–501 (2010).

51. N. W. Moriarty, R. W. Grosse-Kunstleve, P. D. Adams, electronic Ligand Builder and Optimization Workbench (eLBOW): a tool for ligand coordinate and restraint generation. Acta Crystallogr. D Biol. Crystallogr. 65, 1074–1080 (2009).

52. The PyMOL Molecular Graphics System, Version 1.5.0.4, Schrödinger, LLC.

