## Supplementary Material for "Inhibition and Crystal Structure of the Human DHTKD1-Thiamin Diphosphate Complex"

**Supplementary Materials**
**Synthesis of phosphonic acids**
**Synthesis of glutarylphosphonic acid.**

**1a.** To a solution of dihydro-2H-pyran-2,6(3H)-dione (3 g, 26.31 mmol, 1 equiv.) and benzyl alcohol (2.85 g, 1 equiv.) in DMF (10 mL) was added with DMAP (0.5 g, 15 mol%), the reaction mixture was stirred at room temperature for 5 h. Then concentrated under high vacuum and purified via FCC (Hexanes/EtOAc, 20% to 50%) to give product 5-(benzyloxy)-5-oxopentanoic acid **1a** (2g, 34%).

**$^1$H-NMR (600 MHz, $CDCl_3$): δ** 7.40-7.34 (m, 5H), 2.47-2.44 (m, 4H), 2.03-1.98 (m, 2H);
**LCMS (TOF-ESI)** for $C_{12}H_{14}O_4$ [M] 222.0892;
**Calculated [M+H]:** 223.0962; **Found [M+ H]$^+$** for 223.0970.

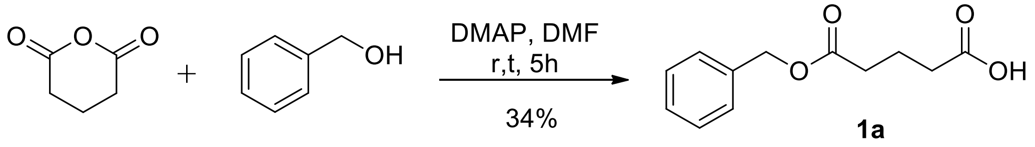

**2a**. To a solution of 5-(benzyloxy)-5-oxopentanoic acid **1a** (1.8 g, 8.1 mmol, 1 equiv.) in $CH_2Cl_2$ (20 mL) was added with $SOCl_2$ (1.64 g, 2.5 equiv.) at room temperature, the whole solution was stirred at reflux for 6 h. After cooled to room temperature, the solution was concentrated to remove any volatiles under high vacuum to give corresponding benzyl 5-chloro-5-oxopentanoate as clear oil which was used for next step without purification.

**$^1$H-NMR (600 MHz, $CDCl_3$): δ** 7.41-7.36 (m, 5 H), 3.02-3.00 (t, 2H), 2.49-2.47 (t, 2H), 2.08-2.03 (m, 2H);

benzyl 5-chloro-5-oxopentanoate (1.0 g, 4.16 mmol, 1 equiv.) was added into trimethyl phosphite (0.57 g, 1.1 equiv.) at 0°C dropwise, upon the completion of addition, the mixture was stirred at room temperature for 12 h. Then the mixture was subjected to high vacuum to remove any unreacted trimethyl phosphite and gave the desired product benzyl 5-(dimethoxyphosphoryl)-5-oxopentanoate **2a** (1.2g, 92%).

**$^1$H-NMR (600 MHz, $CDCl_3$): δ** 7.40-7.28 (m, 5H), 5.14 (s, 2H), 3.88 (s, 3H), 3.86 (s, 3H), 2.95-2.93 (t, 2H), 2.44-2.42 (s, 2H), 2.00-1.97 (m, 2H).

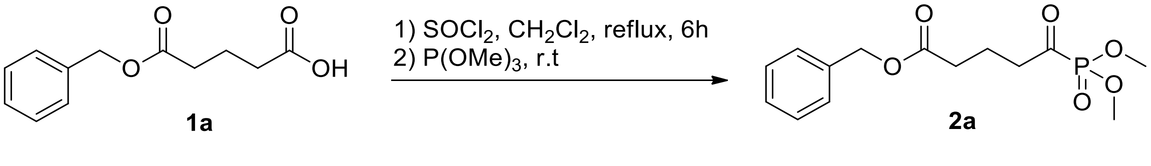

**3a**. To a solution of benzyl 5-(dimethoxyphosphoryl)-5-oxopentanoate **2a** (0.2 g, 0.64 mmol, 1 equiv.) in THF (5 mL) was added with Pd/C (70 mg, 10 mol %), the system was flushed with Argon and hydrogen gas sequentially, and then stirred under $H_2$ (balloon) environment for 12 h. After filtration, the filtrated was collected and concentrated to give product 5-(dimethoxyphosphoryl)-5-oxopentanoic acid as oil.

**$^1$H-NMR (600 MHz, $CDCl_3$): δ** 3.90 (s, 3H), 3.88 (s, 3H), 2.98-2.96 (t, 2H), 2.43- 2.41 (t, 2H), 1.99-1.95 (m, 2H);

To a solution of above obtained 5-(dimethoxyphosphoryl)-5-oxopentanoic acid in $CH_2Cl_2$ (5 mL) was added with TMSBr (0.58 g, 6 equiv.) at 0°C dropwise, upon completion of addition, the reaction was slowly warmed up to room temperature and stirred for 12 h. The reaction was then quenched with $CH_3CN/H_2O$ (1 mL/0.5 mL) and stirred for 1 h. Then solvent was removed and the

residue was washed with $CH_2Cl_2$ and dried under high vacuum to give product 5-oxo-5-phosphonopentanoic acid **3a** as light brown oil.
**$^1$H-NMR (600 MHz, *$d_2$*-$D_2O$): δ** 2.78-2.76 (t, 2H), 2.31-2.25 (tt, 4H), 1.75-1.72 (m, 2H).
**$^{31}$P-NMR(600 MHz, *$d_2$*-$D_2O$): δ:** -2.44.

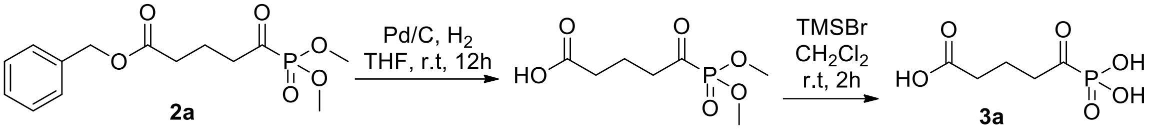

**Synthesis of adipoylphosphonic acid.**
**1b.** A solution of adipic acid (4 g, 2 equiv.), benzyl alcohol **5** (1.5 g, 1 equiv.) and pyridinium p-toluenesulfonate (0.2 g, 10 mol%) in toluene (300 mL) was stirred at reflux for 12h. After cooled to room temperature, the mixture was filtered and the filtrate was collected, concentrated and purified via FCC (Hexanes/EtOAc, 0 to 40%) to give product 6-(benzyloxy)-6-oxohexanoic acid **1b** (0.75 g, 23%) as colorless oil.
**$^1$H-NMR (600 MHz, $CDCl_3$): δ** 7.38-7.35 (m, 5H), 5.14 (s, 2H), 2.43-2.39 (m, 4H), 1.75-1.68 (m, 4H);
**LCMS (TOF-ESI)** for $C_{13}H_{16}O_4$ [M] 236.1049;
**Calculated [M+H]:** 237.1119**; Found [M+ H]$^+$** for 237.1119.

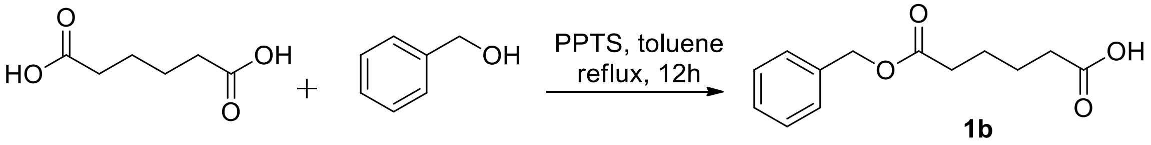

**1bl.** A solution of 6-(benzyloxy)-6-oxohexanoic acid **1b** (0.5 g, 2.12 mmol, 1 equiv.) and DMF (1 drop) in $CH_2Cl_2$ was added with oxalyl chloride (0.54 g, 1.5 equiv.) dropwise at 0°C. The mixture was stirred at room temperature for 2 h and then concentrated under high vacuum to provide product benzyl 6-chloro-6-oxohexanoate as light yellow oil.
**$^1$H-NMR (600 MHz, $CDCl_3$): δ** 7.41-7.35 (m, 5H), 5.14 (s, 2H), 2.94-2.92 (t, 2H), 2.43-2.40 (t, 2H), 1.78-1.72 (m, 4H);

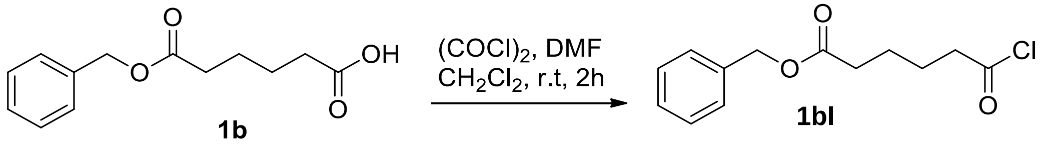

**2b.** benzyl 6-chloro-6-oxohexanoate (0.52 g, 2.05 mmol, 1 equiv.) was added into trimethyl phosphite (0.3 g, 1.1 equiv.) at 0°C dropwise, upon the completion of addition, the mixture was stirred at room temperature for 12 h. Then the mixture was subjected to high vacuum to remove any unreacted trimethyl phosphite and gave the desired product benzyl 6-(dimethoxyphosphoryl)-6-oxohexanoate **2b** (0.64 g, 95%).
**$^1$H-NMR (600 MHz, $CDCl_3$): δ** 7.40-7.34 (m, 5H), 5.13 (s, 2H), 3.88 (d, 3H), 3.87 (d, 3H), 2.88-2.86 (m, 2H), 2.41-2.39 (m, 2H), 1.70-1.66 (m, 4H);
**LCMS (TOF-ESI)** for $C_{15}H_{21}O_6P$ [M] 328.1076;

**Calculated [M+H]:** 329.1146**; Found [M+ H]⁺** for 329.1151.
**$^{31}$P-NMR(600 MHz, *$d_2$*-$D_2O$): δ:** -0.78.

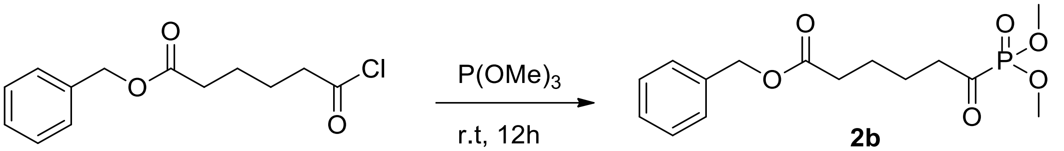

**2b**

**3b.** To a solution of benzyl 6-(dimethoxyphosphoryl)-6-oxohexanoate **2b** (0.2 g, 0.61mmol, 1 equiv.) in THF (5 mL) was added with Pd/C (70 mg, 10 mol %), the system was flushed with Argon and hydrogen gas sequentially, and then stirred under $H_2$ (balloon) environment for 12 h. After filtration, the filtrated was collected and concentrated to give product 6-(dimethoxyphosphoryl)-6-oxohexanoic acid as oil.
To a solution of above obtained 6-(dimethoxyphosphoryl)-6-oxohexanoic acid in $CH_2Cl_2$ (5 mL) was added with TMSBr (0.58 g, 6 equiv.) at 0°C dropwise, upon completion of addition, the reaction was slowly warmed up to room temperature and stirred for 12 h. The reaction was then quenched with $CH_3CN/H_2O$ (1 mL/0.5 mL) and stirred for 1 h. Then solvent was removed and the residue was washed with $CH_2Cl_2$ and dried under high vacuum to give product 6-oxo-6-phosphonohexanoic acid **3b** as resin-like solid.
**$^1$H-NMR (600 MHz, *$d_2$*-$D_2O$): δ** 2.77-2.75(t, 2H), 2.33-2.30 (m, 2H), 1.55-1.49 (m, 4H).
**$^{31}$P-NMR(600 MHz, *$d_2$*-$D_2O$): δ:** -2.25.

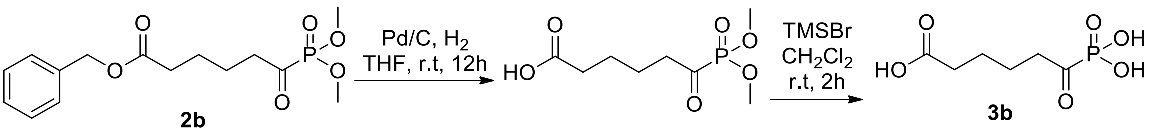

**Synthesis of pimeloylphosphonic acid.**
**1c.** A solution of heptanedioic acid (4.5 g, 2 equiv.), benzyl alcohol (1.5 g, 1 equiv.) and pyridinium p-toluenesulfonate (0.2 g, 10 mol%) in toluene (300 mL) was stirred at reflux for 12h. After cooled to room temperature, the mixture was filtered and the filtrate was collected, concentrated and purified via FCC (Hexanes/EtOAc, 0 to 40%) to give product 7-(benzyloxy)-7-oxoheptanoic acid **1c** (1 g, 29%) as colorless oil.
**$^1$H-NMR (600 MHz, $CDCl_3$): δ** 7.39-7.34 (m, 5H), 5.14 (s, 2H), 2.40-2.35 (m, 4H), 1.72-1.64 (m, 4H), 1.42-1.37 (m, 2H);
**LCMS (TOF-ESI)** for $C_{14}H_{18}O_4$ [M] 250.1205;
**Calculated [M+H]:** 251.1275**; Found [M+ H]⁺** for 251.1284.

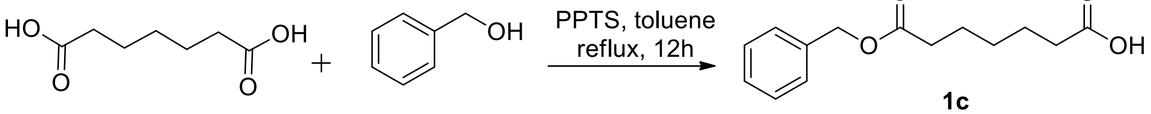

**1c**

**1cl**. A solution of 6-(benzyloxy)-6-oxohexanoic acid **1c** (0.54 g, 2.12 mmol, 1 equiv.) and DMF (1 drop) in $CH_2Cl_2$ was added with oxalyl chloride (0.54 g, 1.5 equiv.) dropwise at 0°C. The mixture

was stirred at room temperature for 2h and then concentrated under high vacuum to provide product benzyl 7-chloro-7-oxoheptanoate as light yellow oil.
**$^1$H-NMR (600 MHz, $CDCl_3$): δ** 7.40-7.35 (m, 5H), 5.14 (s, 2H), 2.91-2.88 (t, 2H), 2.40-2.38 (t, 2H), 1.76-1.72 (q, 2H), 1.71-1.67 (q, 2H), 1.43-1.38 (m, 2H).

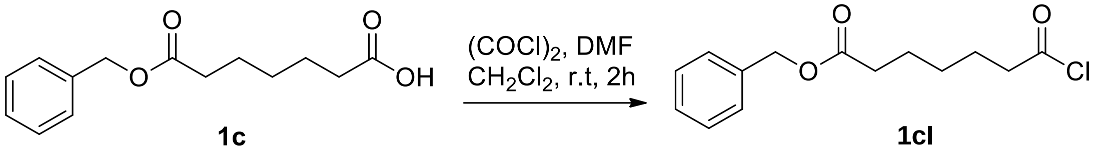

**1c** **1cl**

**2c**. benzyl 7-chloro-7-oxoheptanoate (0.57 g, 2.05 mmol, 1 equiv.) was added into trimethyl phosphite (0.3 g, 1.1 equiv.) at 0°C dropwise, upon the completion of addition, the mixture was stirred at room temperature for 12 h. Then the mixture was subjected to high vacuum to remove any unreacted trimethyl phosphite and gave the desired product benzyl 7-(dimethoxyphosphoryl)-7-oxoheptanoate **2c** (0.7 g, 95%).
**$^1$H-NMR (600 MHz, $CDCl_3$): δ** 7.40-7.34 (m, 5H), 5.13 (s, 2H), 3.89 (s, 3H), 3.87 (s, 3H), 2.85-2.83 (t, 2H), 2.39-2.36 (t, 2H), 1.70-1.64 (m, 4H), 1.38-1.35 (m, 2H);
**LCMS (TOF-ESI)** for $C_{16}H_{23}O_6P$ [M] 342.1232;
**Calculated [M+H]:** 343.1302**; Found [M+ H]$^+$** for 343.1302.
**$^{31}$P-NMR(600 MHz, *d*$_2$-$D_2O$): δ:** -0.68.

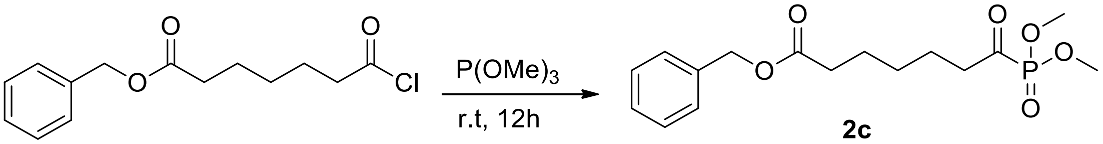

**2c**

**3c**. To a solution of benzyl benzyl 7-(dimethoxyphosphoryl)-7-oxoheptanoate **2c** (0.21 g, 0.61 mmol, 1 equiv.) in THF (5 mL) was added with Pd/C (70 mg, 10 mol %), the system was flushed with Argon and hydrogen gas sequentially, and then stirred under $H_2$ (balloon) environment for 12 h. After filtration, the filtrated was collected and concentrated to give product 7-(dimethoxyphosphoryl)-7-oxoheptanoic acid as oil.
To a solution of above obtained 7-(dimethoxyphosphoryl)-7-oxoheptanoic acid in $CH_2Cl_2$ (5 mL) was added with TMSBr (0.58 g, 6 equiv.) at 0°C dropwise, upon completion of addition, the reaction was slowly warmed up to room temperature and stirred for 12 h. The reaction was then quenched with $CH_3CN/H_2O$ (1 mL/0.5 mL) and stirred for 1 h. Then solvent was removed and the residue was washed with $CH_2Cl_2$ and dried under high vacuum to give product 7-oxo-7-phosphonoheptanoic acid **3c** as white solid.
**$^1$H-NMR (600 MHz, $CDCl_3$): δ** 2.76-2.73 (t, 2H), 2.30-2.28 (m, 2H), 1.53-1.49 (m, 4H), 1.27-1.23 (m, 2H);
**$^{31}$P-NMR(600 MHz, *d*$_2$-$D_2O$): δ:** -2.18.

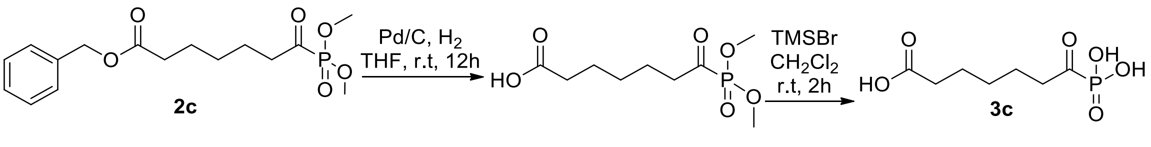

**2c** **3c**

**Repurchase of HTS compounds of interest.** After dose-response evaluation of HTS hits, 13 out 17 hit compounds were commercially available and were purchased for further confirmation in DHTKD1 inhibition assays. HTS ID 255785 (Omeprazole, #K253), Esomeprazole (#J91970), and Omeprazole Sulfone (#4281AC) were purchased from AK Scientific. HTS ID 255785 (Omeprazole, #458520010), HTS ID 285797 (Pantoprazole sodium salt hydrate, #460870010) were obtained from Acros Organics. HTS ID 757002 (Tenatoprazole, #B1848) was from APExBio. HTS ID 130002 (Bithionol, #T0865) and Pantoprazole Sulfide (#P2066) were purchased from TCI America. HTS ID 48508 (#5563352), HTS ID 49582 (#5676527) were obtained from ChemBridge. HTS ID 155744 (#5586-4259) and HTS ID 45447 (#R052-1687) were from ChemDiv. HTS ID 29708 (#Z44101855) was obtained from Enamine. HTS ID 49098 (#STK758032), HTS ID 49259 (#STL323879), HTS ID 48303 (#STK398483) and HTS ID 36573 (#STL297498) were purchased from Vitas-M Laboratory.

### Supplementary Figures

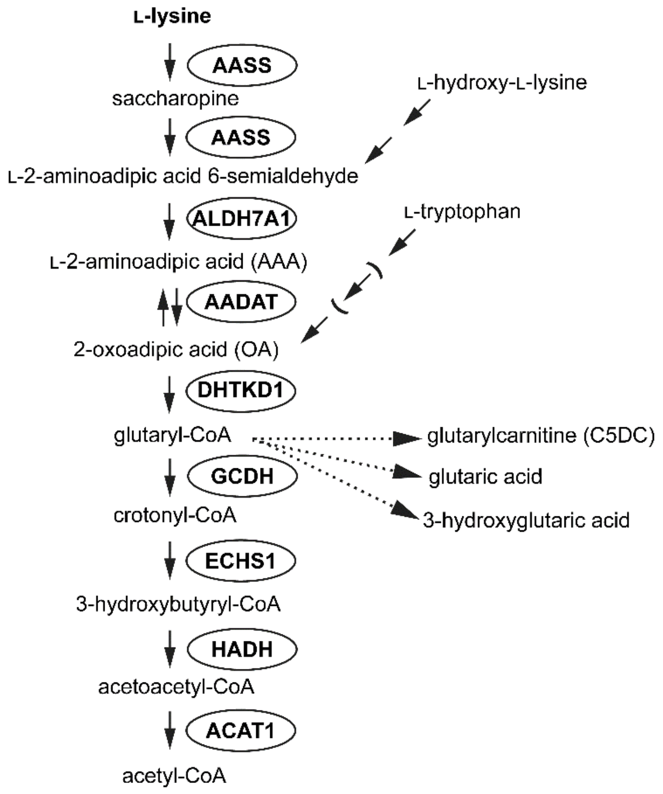

**Figure S1.** The degradation of L-lysine occurs mainly via the mitochondrial saccharopine pathway. The pathway consists of nine different enzymatic steps that ultimately yield 2 acetyl-CoAs and several reducing equivalents. Degradation of L-hydroxy-L-lysine leads to L-2-aminoadipic acid 6-semialdehyde and cytosolic degradation of tryptophan with kynurenine as intermediate leads to the production of 2-oxoadipic acid (OA). The intermediate OA is further metabolized into glutaryl-CoA via oxidative decarboxylation by the DHTKD1, the E1 component of the 2-oxoadipic acid dehydrogenase complex (OADHc). Defective DHTKD1 leads to accumulation of OA and AAA (the level of AAA correlates with the amount of OA by reversible transamination catalyzed by AADAT). Glutaryl-CoA is converted into crotonyl-CoA via oxidative decarboxylation by GCDH. The GCDH protein is deficient in GA1 leading to the accumulation of glutaryl-CoA metabolites: glutarylcarnitine (C5DC), glutaric acid and 3-hydroxyglutaric acid. AASS, alpha-aminoadipic semialdehyde synthase; ALDH7A1, alpha-aminoadipic semialdehyde dehydrogenase; AADAT, kynurenine/alpha-aminoadipate aminotransferase; DHTKD1, dehydrogenase E1 and transketolase domain containing 1; GCDH, glutaryl-CoA dehydrogenase; ECHS1, enoyl-CoA hydratase; HADH, hydroxyacyl-coenzyme A dehydrogenase; ACAT1, acetyl-CoA acetyltransferase.

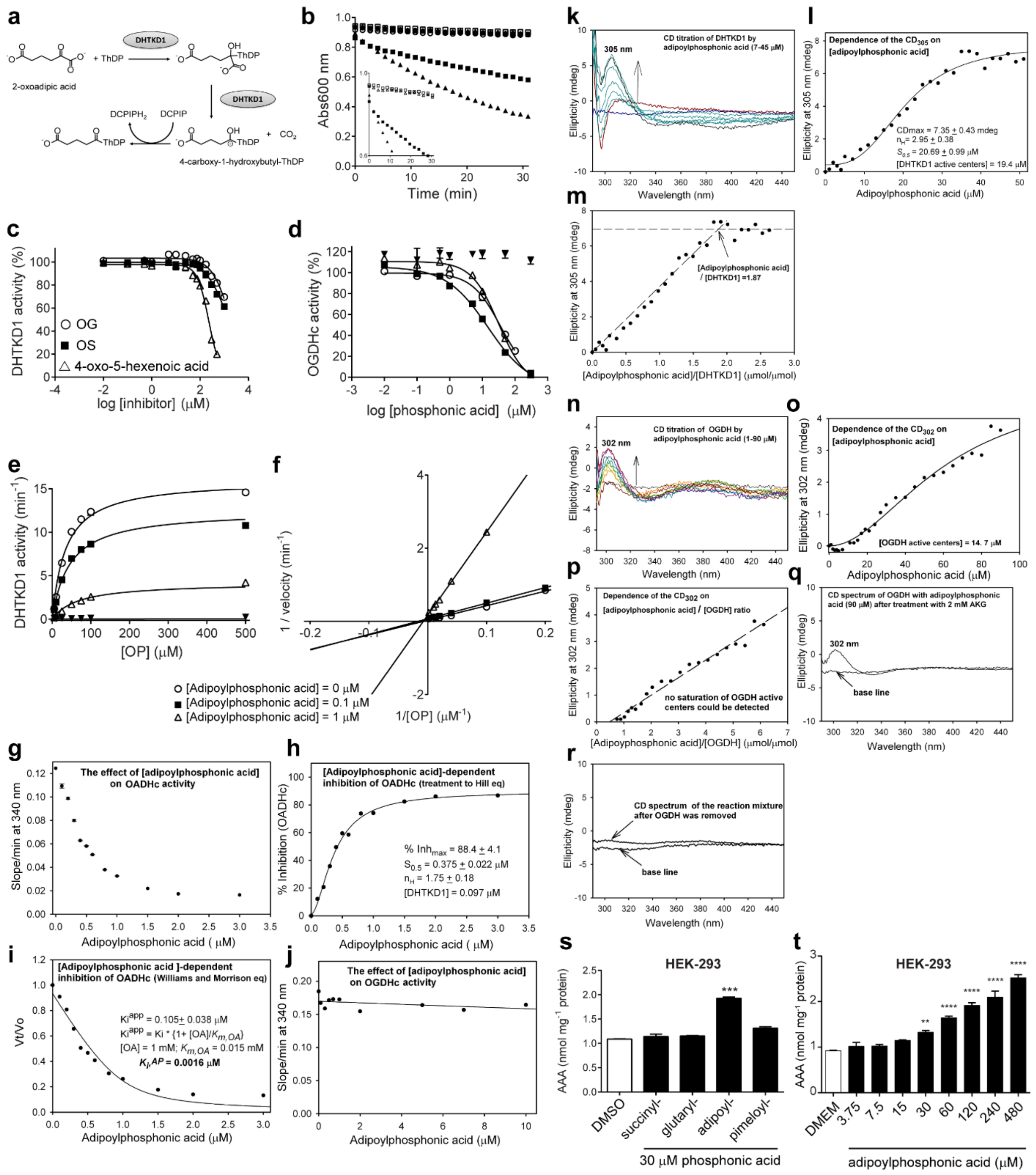

**Figure S2.** DHTKD1 substrate analog inhibitors. (a) DHTKD1 reaction using dark blue DCPIP as an artificial electron acceptor that turns transparent after reduction (DCPIPH$_2$). (b) Reaction progress curves of DHTKD1 using 62.5 μM OG (●), 62.5 μM OA (■) and 62.5 μM OP (▲) as substrates. Open symbols are the corresponding blanks. The inset (left-hand side) shows a close-up of the OA and OP curves at the start of the reaction. The DCPIP-based assay for E1 component was used, where DCPIP acts as an electron acceptor. The reaction was monitored by the decrease of absorbance at 600nm corresponding to the formation of DCPIPH$_2$. (c) OG and OS are weak inhibitors of DHTKD1 while 4-oxo-5-hexenoic acid is an irreversible inhibitor. Preincubation of DHTKD1 with 4-oxo-5-hexenoic acid leads to a time-dependent increase in inhibition and a mass shift is observed upon separation in a SDS-PAGE gel, pointing to a covalent modification of the enzyme (data not shown). Dose-response curves of DHTKD1 inhibition by OG (○), OS (■) and by 4-oxo-5-hexenoic acid (△), using OP as substrate. The $IC_{50}$ of OG, OS and 4-oxo-5-hexenoic acid are 1767 ± 308, 1470 ± 139 and 229 ± 10 μM, respectively. (d) Dose-response curves of porcine heart OGDHc inhibition by succinyl- (○), glutaryl- (■), adipoyl- (△) and pimeloyl- (▼) phosphonic acids, using OG as substrate. (e and f) Adipoylphosphonic acid is a DHTKD1 mixed competitive-noncompetitive inhibitor. DHTKD1 activity was measured at different concentrations of adipoylphosphonic acid: 0 (○), 0.1 (■), 1 (△) and 300 μM (▼), in the presence of 62.5 μM OP as substrate. A mixed-model of inhibition was used with the following parameters: $V_{max}$ = 16.05 ± 0.46 $min^{-1}$, $K_m$ = 34.9 ± 3.0 μM, $K_i$ = 0.15 ± 0.03 μM and α = 2.3 ± 0.8 (e). Double reciprocal plot showing inhibition of DHTKD1 by adipoylphosphonic acid. Linear regression analysis is consistent with mixed competitive-noncompetitive type of inhibition (f). (g–j) Inhibition of the human OADHc and OGDHc by adipoylphosphonic acid. Plot of the reaction rates as a function of [adipoylphosphonic acid]. The OADHc was assembled from DHTKD1 (0.4 mg), DLST (0.80 mg) and DLD (5.0 mg) components in 0.1 M Tris.HCl (pH 7.5) containing 0.3 M $NH_4Cl$, 0.5 mM ThDP and 2.0 mM $MgCl_2$. After 40 min of incubation at 25 °C, an aliquot containing 0.01 mg of DHTKD1 and the corresponding amounts of DLST and DLD was withdrawn and was placed into 1 ml of the reaction assay containing all components necessary for the overall activity assay in the absence (control) and in the presence of 0.1-3 μM adipoylphosphonic acid. After 10 min incubation at 25 °C, the overall activity measurement was initiated by addition of 2-oxoadipic acid (1.0 mM) and CoA (400 μM) and the progress curves were recorded at 37 °C for 3 min. The reaction rates (slope/min at 340 nm) were calculated from a linear part of the recorded progress curves and were plotted versus [adipoylphosphonic acid] (g). Plot of % inhibition as a function of [adipoylphosphonic acid]. The solid curve shows the least-squares fit of the data to the Hill plot equation: $y = yo + (V_{max} * [\text{adipoylphosphonic acid}]^n)/(S_{0.5}{}^n + [\text{adipoylphosphonic acid}]^n)$ (h). Plot of steady-state fractional velocity as a function of [adipoylphosphonic acid]. The solid curve shows the least-squares fit of the data to the Morrison equation for tight-binding inhibitors: $v/v_o = \{[E] - [I] - K_i^{app} + \text{sqrt}([E] – [I] –K_i^{app})^2 + 4[E]^* K_i^{app}\}/2E$, where [E] is the concentration of DHTKD1 active centers, [I] is the total concentration of adipoylphosphonic acid, $K_i^{app}$ is the dissociation constant of adipoylphosphonic acid, v is the initial reaction velocity at different concentrations of the adipoylphosphonic acid, $v_o$, is the initial velocity in the absence of adipoylpohosphonic acid. If the inhibitor and substrate compete for the same active site, $K_i^{app}$ is related to the dissociation constant of the inhibitor ($K_i$) by the following equation: $K_i^{app} = K_i\ (1+[S]/K_m)$ where $K_m$ is the Michaelis constant and [S] is the concentration of substrate (i). Plot of the reaction rates as a function of [adipoylphosphonic acid] for OGDHc with 2-oxoglutaric acid as substrate. The OGDH was assembled with DLST and DLD into OGDHc at 25 °C for 40 min similarly to that for DHTKD1 above. The overall activity measurement was initiated by addition of 2-oxoglutaric acid (2.0 mM) and CoA (400 μM) and the progress curves were recorded at 37 °C for 1 min (j). (k–m) Circular dichroism titration of DHTKD1 by adipoylphosphonic acid. The DHTKD1 (19.4 μM concentration of active centers) in 100 mM HEPES (pH 7.5) containing 0.5 mM ThDP, 2.0 mM $MgCl_2$, 0.15 M NaCl and 5% glycerol was titrated by adipoylphosphonic acid (1-51 μM) (k). The intensity of the CD band at 305 nm was estimated and was plotted versus the concentration of adipoylphosphonic acid (l) or versus the [adipoylphosphonic acid]/[DHTKD1 active centers] ratio (m). (n–r) Circular dichroism titration of human OGDH by adipoylphosphonic acid. The OGDH (concentration of active centers of 14.7 μM) in 100 mM HEPES (pH 7.5) containing 0.5 mM ThDP, 2.0 mM $MgCl_2$, 0.15 M NaCl and 5% glycerol was titrated by adipoylphosphonic acid (1-80 μM) (n). The intensity of the CD band at 302 nm was estimated and was plotted versus the concentration of adipoylphosphonic acid (o) or versus the [adipoylphosphonic acid]/[OGDH active centers] ratio (p). The CD band at 302 nm was still observed after overnight incubation with 2.0 mM 2-

oxoglutaric acid; however, its intensity was reduced indicating weak binding of adipoylphosphonic acid to the active centers of OGDH and/or its partial replacement by an excess of 2-oxoglutaric acid overnight (q). On separation of OGDH from the reaction mixture, no CD band at 302 nm was detected as shown in panel (r), clearly demonstrating that the observed $CD_{302}$ is a signature of the DHTKD1-bound pre-decarboxylation intermediate rather than product formation. (s and t) Adipoylphosphonic acid inhibits DHTKD1 in a cellular model. HEK-293 cellular AAA levels in the presence of the phosphonic acids in DMEM medium (0.8 mM L-lysine) (s) and AAA levels with increasing concentrations of the DHTKD1 inhibitor adipoylphosphonic acid in DMEM medium (0.8 mM L-lysine). **, $P < 0.01$; ***, $P < 0.001$; ****, $P < 0.0001$.

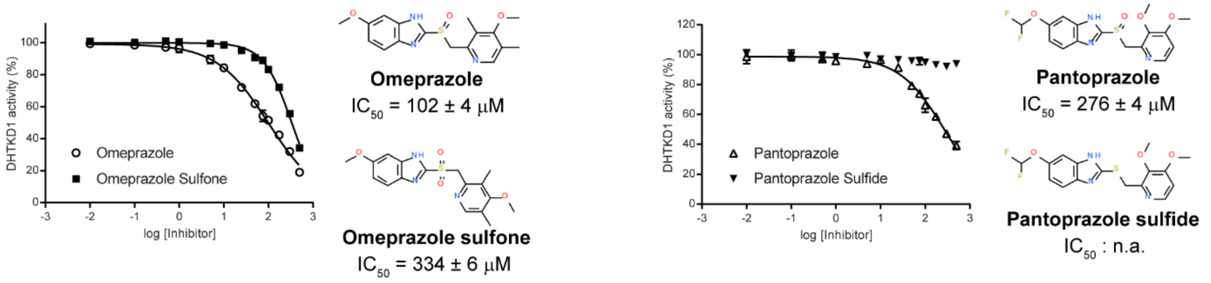

**Figure S3.** Dose-response curves of omeprazole and pantoprazole and their sulfone and sulfide analogs, respectively.

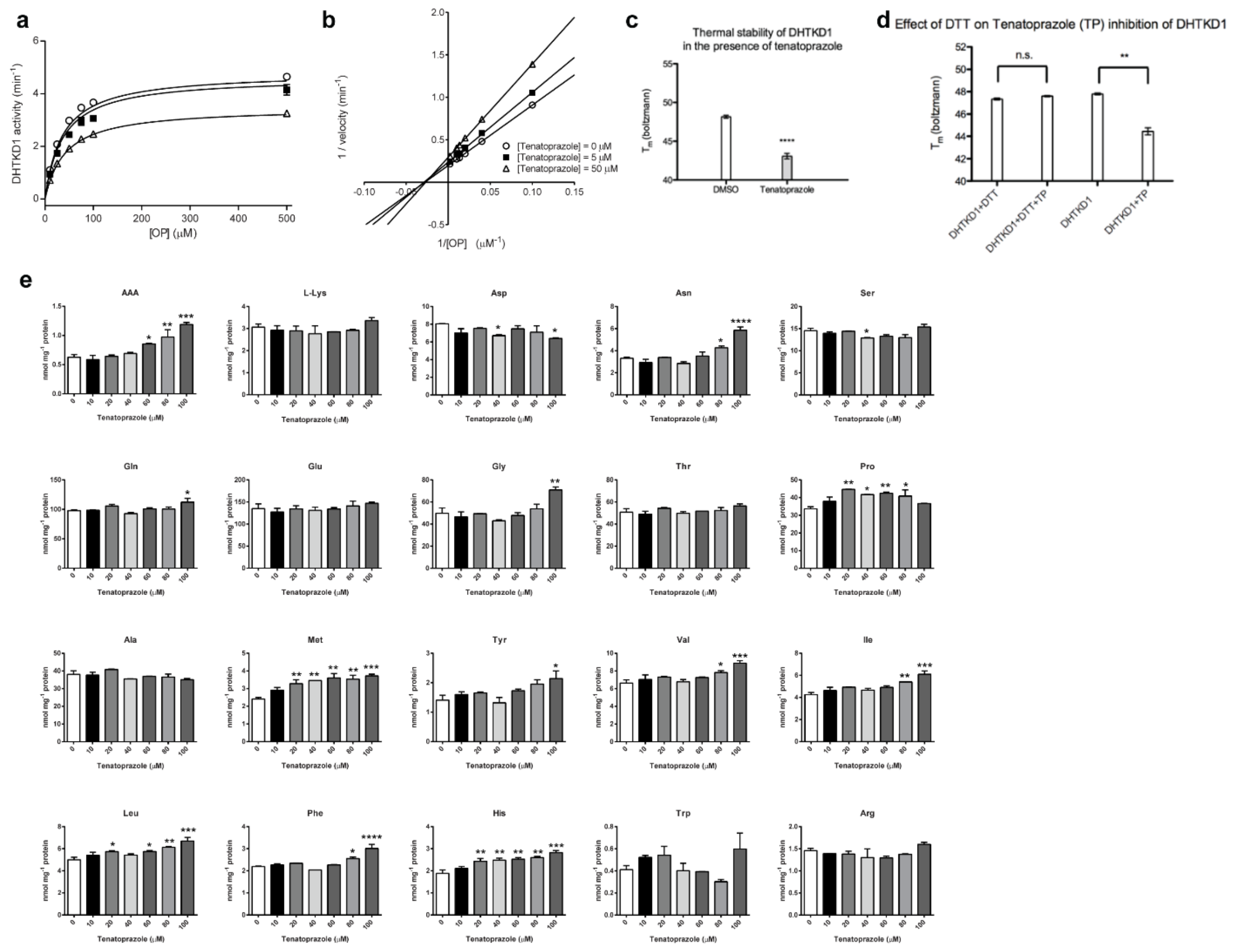

**Figure S4.** Tenatoprazole is a noncompetitive DHTKD1 inhibitor. (a) DHTKD1 was incubated with 0 (○), 5 (■) and 50 μM (△) of tenatoprazole. A mixed model of inhibition was used with $V_{max}$ = 4.8 ± 0.1 min$^{-1}$, $K_m$ = 36 ± 4 μM, $K_i$ = 83 ± 34 μM and $\alpha$ = 1.5 ± 0.9. (b) Double reciprocal plot showing inhibition of DHTKD1 by tenatoprazole. Linear regression analysis is consistent with a noncompetitive type of inhibition. (c) Thermal stability of DHTKD1 with or without 500 μM tenatoprazole. (d) Effect of DTT on tenatoprazole thermal destabilization of DHTKD1. Tenatoprazole at 200 μM or DMSO was incubated with DHTKD1 in the presence or absence of 2 mM DTT. (e) Tenatoprazole was incubated with increasing concentrations of tenatoprazole and the level of intracellular amino acids was evaluated. Tenatoprazole was toxic to the cells above 100 μM. *, $P < 0.05$; **, $P < 0.01$; ***, $P < 0.001$; ****, $P < 0.0001$.

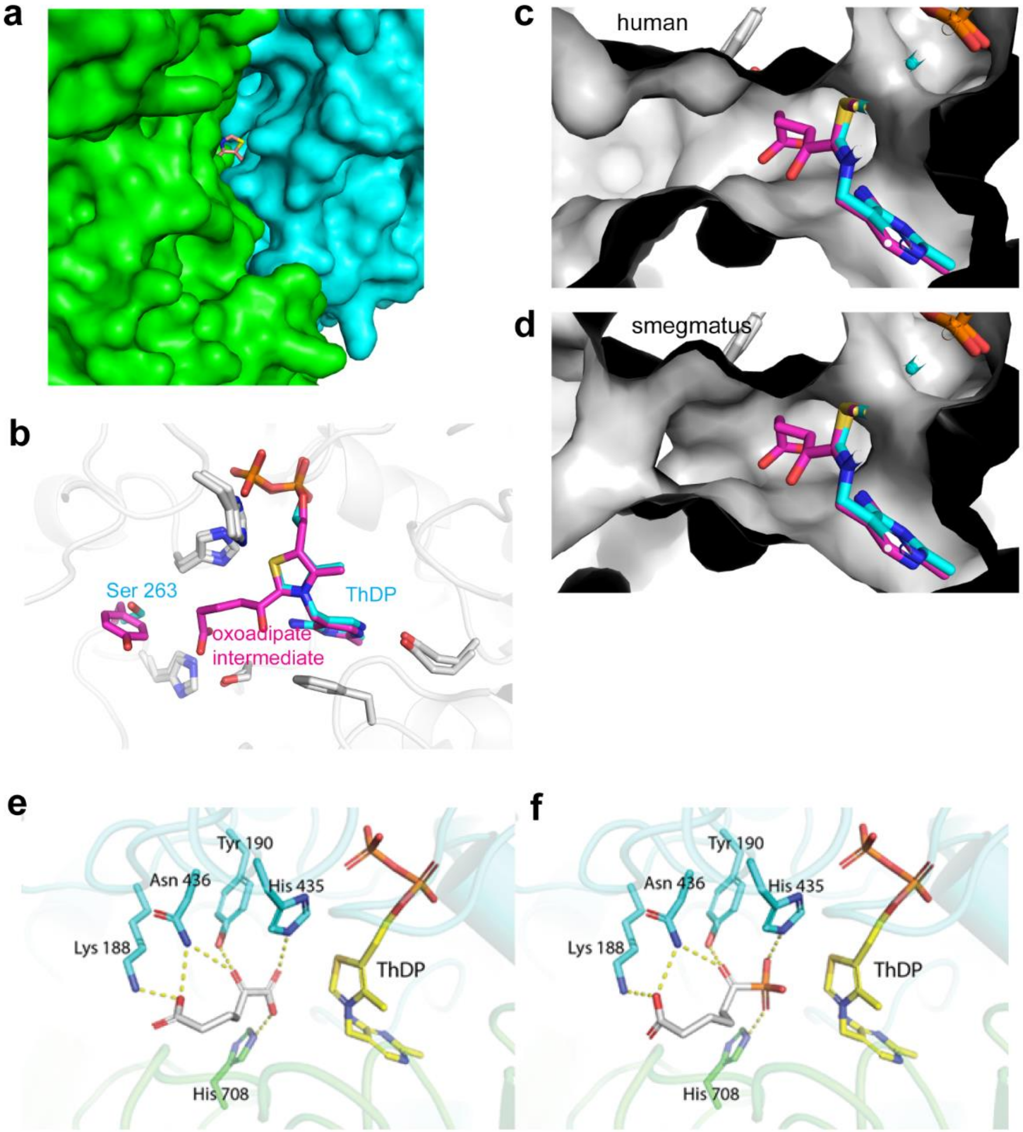

**Figure S5.** DHTKD1 structure, comparison with *M. smegmatis* KGD and docking of the substrate OA and the inhibitor adipoylphosphonic acid. (a) View of the ThDP channel from outside the protein. (b) Overlay of active sites of *M. smegmatis* (PDB code 3ZHU, (1)) and our human DHTKD1 structure. (c) Surface rendering of human active site with modeled in oxoadipate intermediate. (d) Surface rendering of *M. smegmatis* structure with the OA intermediate. (e and f) Docking complex of OA (e) and docking complex of adipoylphosphonic acid (f). Polar contacts (hydrogen bonds and ionic interactions) interactions are shown as yellow dashed lines. Residues are colored according to the chain they belong to.

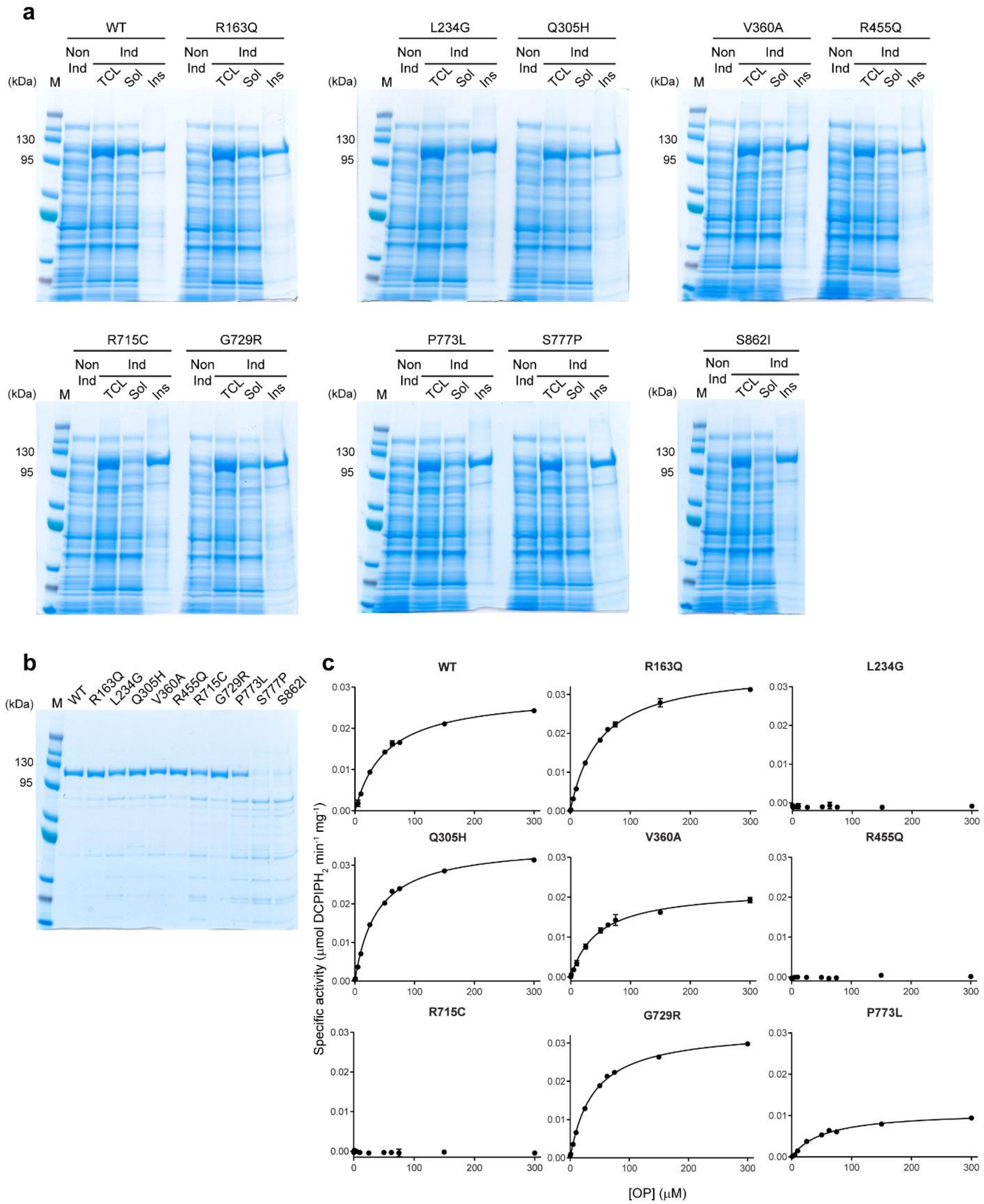

**Figure S6.** DHTKD1 variant expression and enzyme kinetics. (a) SDS-PAGE analysis of the solubility of the different DHTKD1 variants. M, low molecular mass marker; Non Ind, total cell lysate before induction; Ind, IPTG induced fraction; TCL, total cell lysate after 24 h induction of recombinant human DHTKD1 by 1 mM IPTG at RT; Sol, soluble fraction; Ins, insoluble fraction. (b) Analysis of purified DHTKD1 (wild-type and variants) by SDS-PAGE. (c) The effect of substrate

(OP) concentration on the catalytic activity of DHTKD1 wild type and variants. The DHTKD1 activity was assayed at standard conditions (0 – 300 μM OP, 2 mM $MgCl_2$, 1 mM ThDP and 30 °C). The data was analyzed by non-linear regression analysis using the GraphPad Prism 7 software and the Michaelis-Menten equation. The kinetic constants are summarized in Table 1.

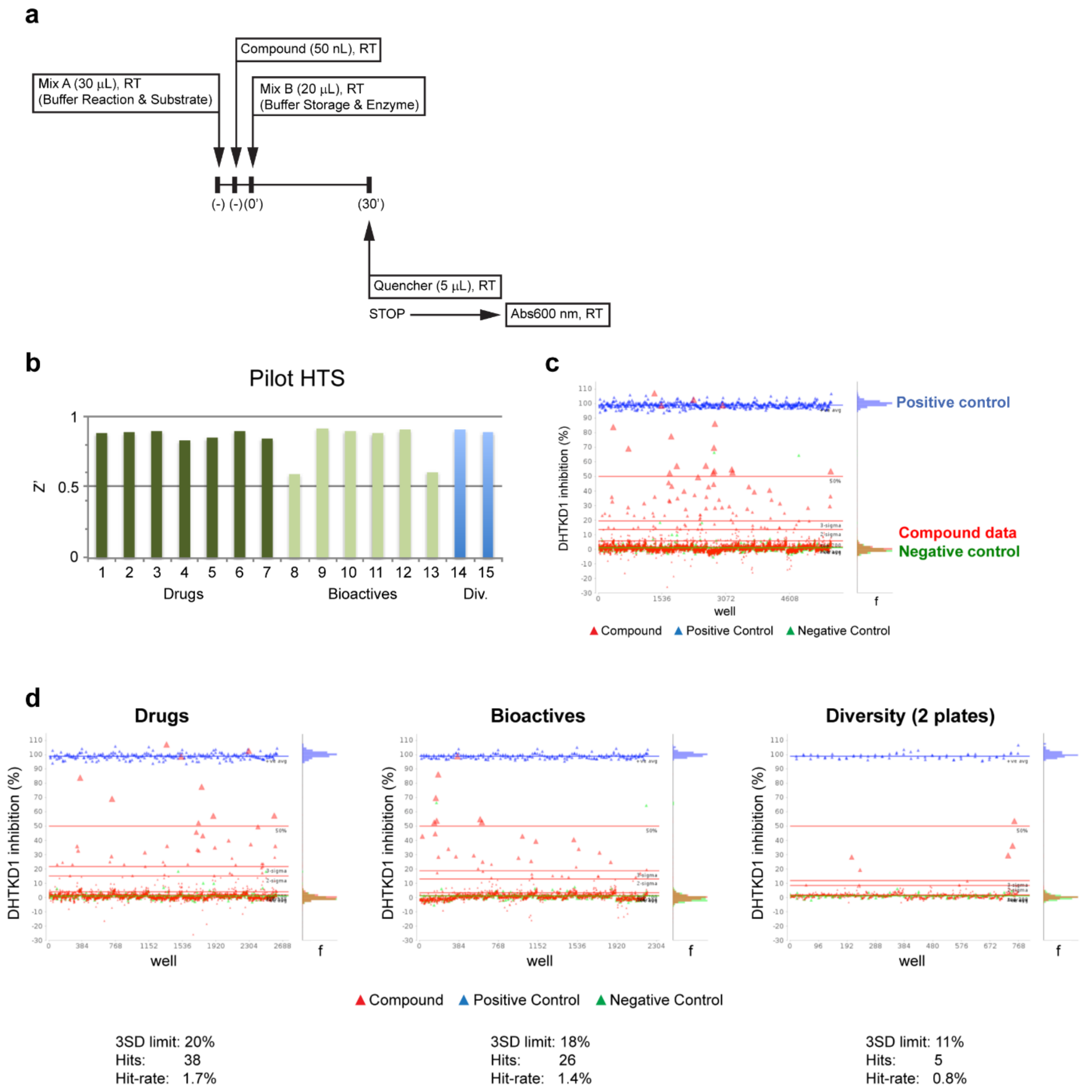

**Figure S7.** Pilot HTS and assay quality indicators for identification of DHTKD1 inhibitors. (a) Schematic illustration of the inhibitor screen. Mix A (Buffer reaction & Substrate, final concentration): 50 mM MOPS, pH 7.4, 100 mM KCl, 2 mM $MgCl_2$, 1 mM ThDP, 0.1 mM DCPIP, 0.01% Triton X-100, 0.25% Prionex and 62.5 µM OP; Mix B (Buffer storage & Enzyme, final concentration): 1.25 µg DHTKD1 in dialysis buffer; Quencher: 100 µM $ZnCl_2$, 0.01% Triton X-100. (b) Z' performance over the 15 assay plates included in pilot phase: 2,177 compounds from the Drug collection (Pharmakon and SMDC drugs), 1,835 compounds from the Bioactive collection (SelleckChem bioactive) and 2 plates from the diversity collection (ChemBridge). The Z' was ~0.8 (a Z′ of 0.5 to 1.0 indicates a significant separation between positive and negative controls). The percent coefficient of variance (% CV) was < 5% (a % CV of <15% is considered acceptable). (c) Scatter plot of DHTKD1 percentage inhibition of each well is shown for the screen pilot phase. Each symbol represents an assay well. Blue line is the mean inhibition of positive control (adipoylphosphonic acid inhibitor); red lines represent '50% inhibition', '3-sigma', and '2-sigma'

above the mean of the population of compounds tested. (d) Shows the same data as in (c), but the compounds tested have been grouped by library collection: drugs, bioactives and 2 plates from the diversity collection. The percentage of active compounds, number of hits, and hit-rate above three standard deviations are indicated for each collection. For each scatter plot the blue line is the mean inhibition of positive control (adipoylphosphonic acid); red lines represent ‘50% inhibition’, ‘3-sigma’, and ‘2-sigma’ above the mean of the population of compounds tested. Green line represents the mean inhibition of negative control.

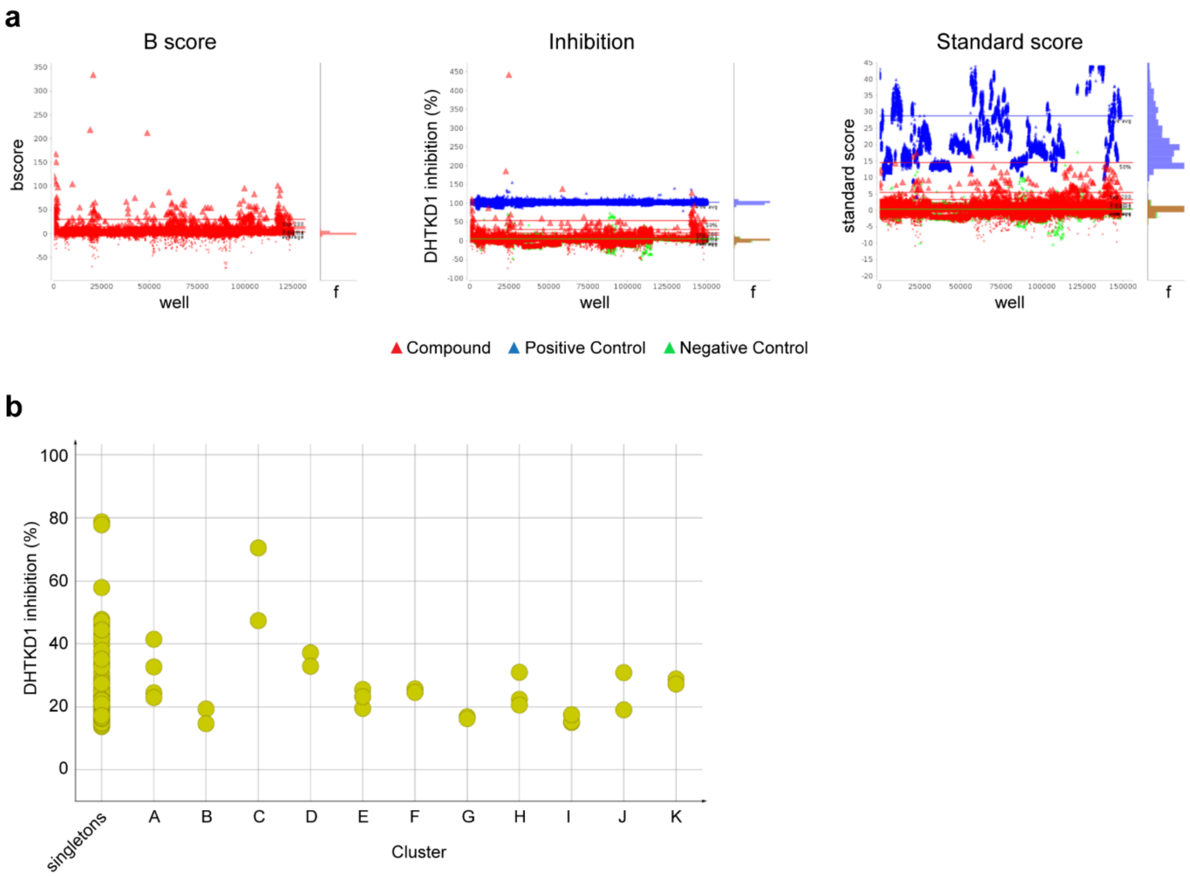

**Figure S8.** HTS for identification of DHTKD1 inhibitors. (a) The B score, the DHTKD1 inhibition and standard score by well are shown for the HTS. Each symbol represents an assay well. The hit selection threshold was set at calculate mean + 3SD for control dependent (% inhibition) and independent (B score and standard score) activity parameters. The candidates must satisfy statistical criteria for all three parameters. Some outlier data points (inhibition ≥100%) are occasionally seen. Blue line +ve avg, mean inhibition of positive control (adipoylphosphonic acid); red lines '3-sigma', '50% inhibition', '2-sigma' above the mean of the population of compounds tested; red line top 200, top 200 active compounds above the mean of the population of compounds tested; red line average, mean of the population of compounds tested. Green line average, mean inhibition of negative control. Right graph represents the histogram of the compounds tested. (b) HTS hit compounds meeting threshold criteria and after exclusion of persistent high absorbance row effect. 133 hit compounds were identified, including 106 singletons and 27 compounds in 11 clusters (≥2 members, A to K). Hit data clustered in DataWarrior, using default similarity limit = 0.8.

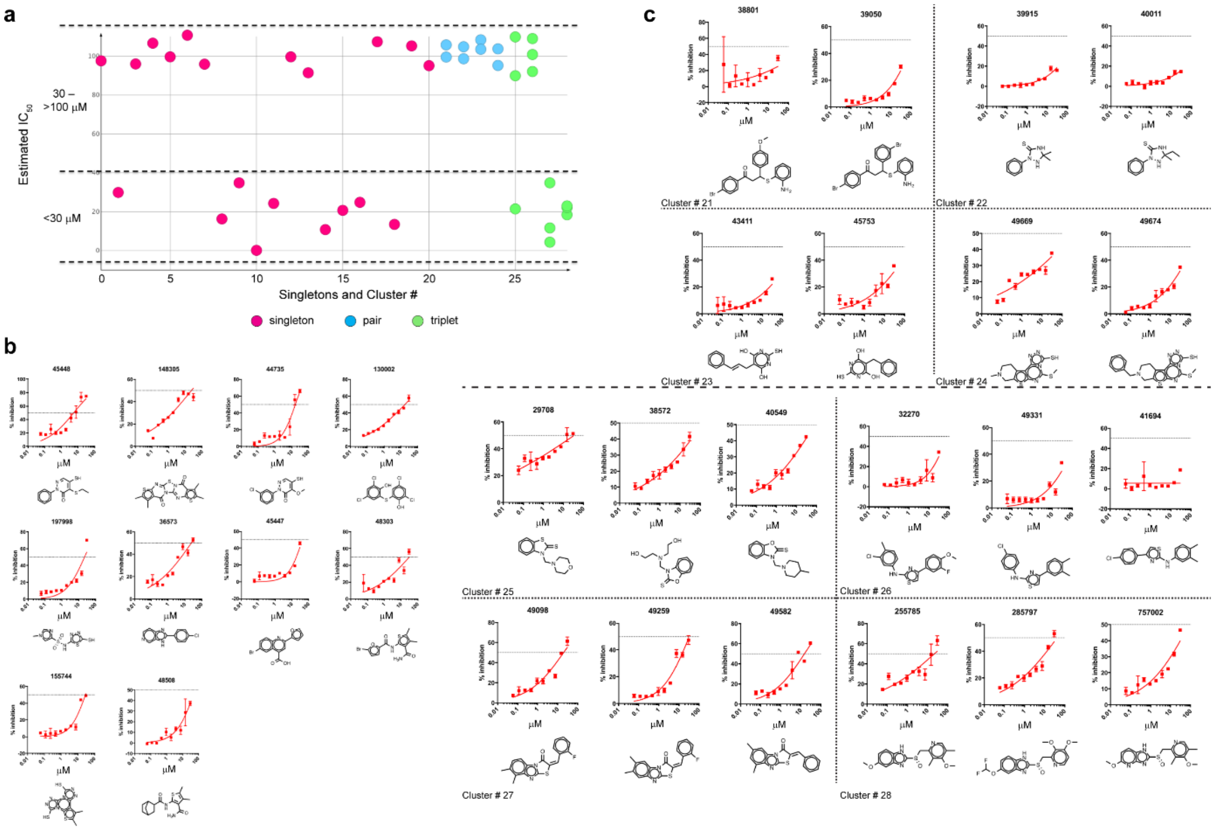

**Figure S9.** DHTKD1 inhibition concentration-response curves. (a) 40 out of 192 HTS hit compounds exhibited concentration responsive inhibition. 16 hit compounds had estimated $IC_{50}$ ≤ 30 μM (9 singletons and 7 compounds from 3 clusters). (b) Chemical structures and dose-response curves for singletons with $IC_{50}$ ≤ 30 μM. After evaluation of 64 extra HTS hit compounds, the hit 48508 ($IC_{50}$ ~50 μM) was added to the singleton list. (c) Chemical structures and dose-response curves for compounds belonging to 8 structure clusters. No molecules retested yield complete dose curves, with greatest inhibition seen at ~75%. Several compounds display rollover (apparent saturation) at moderate inhibition, possibly due to poor compound solubility.

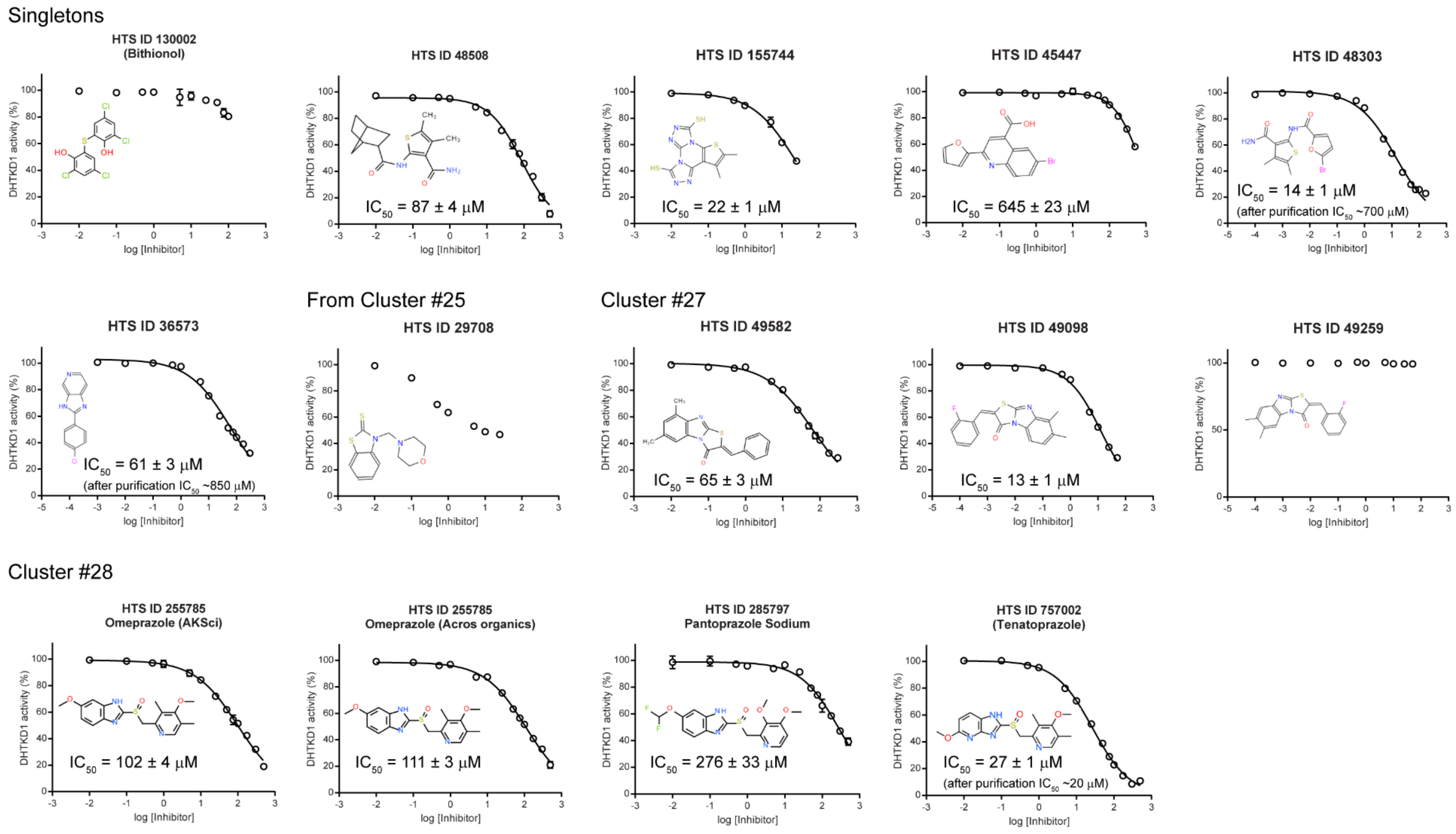

**Figure S10.** Confirmation of DHTKD1 HTS inhibitors. Chemical structures and dose-response curves of 13 repurchased hit compounds from commercial sources for confirmation. Hits 48303, 36573 and 757002 were tested as repurchased material and after re-purification. The first two displayed much higher $IC_{50}$s values after purification.

### Supplementary Tables

**Table S1. Evaluation of DHTKD1 substrate analogs as inhibitors and confirmed HTS hits.**

| Compound | Inhibition[a,b] (at 300 μM) (%) | $IC_{50}$ (μM) | $K_i$ (μM) | Structure |
|---|---|---|---|---|
| 2-oxoglutaric acid (OG)[c] | 12.4 ± 0.6<br>18.2 ± 0.5 (500 μM)<br>30.4 ± 0.8 (1000 μM) | 1767 ± 308 | 250 (competitive) | 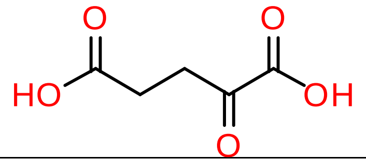 |
| 2-oxosuberic acid (OS)[c] | 13.3 ± 1.1 | 1470 ± 139 | (likely competitive) | 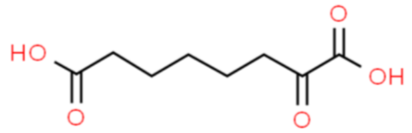 |
| 4-oxo-pimelic acid | 0 | – | – | 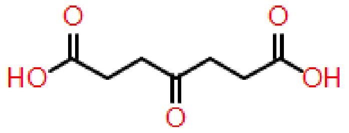 |
| adipic acid[c] | 0.9 ± 3.1 | – | – | 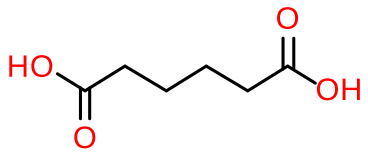 |
| vigabatrin[c] (4-amino-5-hexenoic acid) | -0.5 ± 0.6 | – | – | 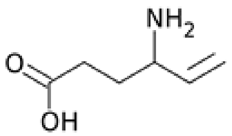 |
| 4-oxo-5-hexenoic acid | 67.4 ± 0.4<br>80.1 ± 0.3 (500 μM) | 229 ± 10 | 267 (irreversible) | 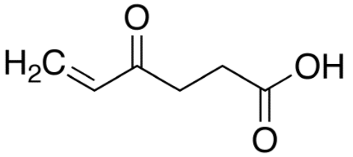 |

| | | | | |
|---|---|---|---|---|
| succinylphosphonic acid (C4) | 1.3 ± 0.3 (100 μM) | – | – | 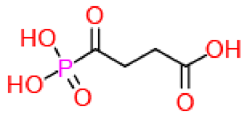 |
| glutarylphosphonic acid (C5) | 59.2 ± 1.7 (100 μM)<br>79.7 ± 0.2 | 55.1 ± 2.3 | n.d.[d] | 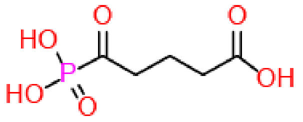 |
| adipoylphosphonic acid (C6) | 98.4 ± 0.4 (100 μM)<br>100.4 ± 0.4 | 0.21 ± 0.02 | 0.15 ± 0.03 (mixed competitive-noncompetitive inhibitor) | 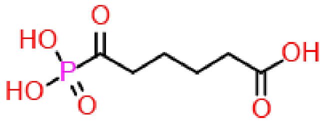 |
| pimeloylphosphonic acid (C7) | 43.8 ± 4.7 (100 μM)<br>74.9 ± 2.9 | 73.5 ± 3.1 | n.d. | 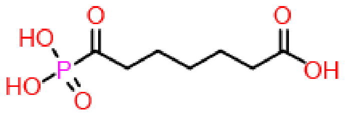 |
| 5-phosphonopentanoic acid | 1.6 ± 1.6 | – | – | 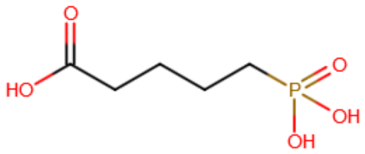 |
| dodecanoic acid | 7.2 ± 2.0 | – | – | 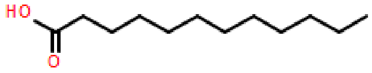 |
| dodecanedioic acid | 3.9 ± 1.2 | – | – | 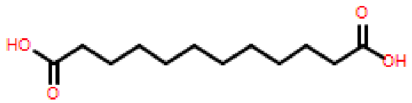 |

| | | | | |
|---|---|---|---|---|
| suberic acid | -2.1 ± 2.8 | – | – | 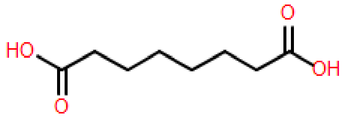 |
| sebacic acid | 3.5 ± 2.4 | – | – | 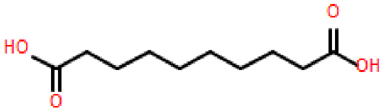 |
| tetradecanedioic acid | 2.8 ± 0.2 | – | – | 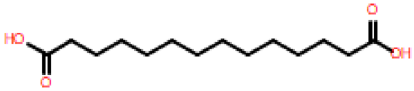 |
| 2-ketoisocaproic acid | 2.8 ± 0.7 | – | – | 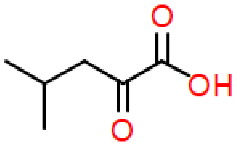 |
| 6-oxohexanoic acid | 2.7 ± 1.5<br>8.5 ± 2.4 (500 μM) | – | – | 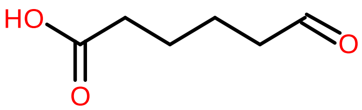 |
| metformin[c] | 0.4 ± 1.5 | – | – | 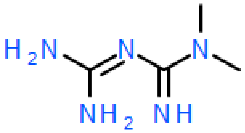 |
| N-Oxalylglycine[c] | -4.3 ± 3.6 | | | 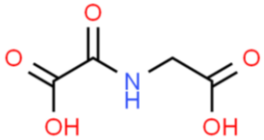 |

| | | | | |
|---|---|---|---|---|
| 2-ketocaproic acid | 1.5 ± 2.3 | – | – | 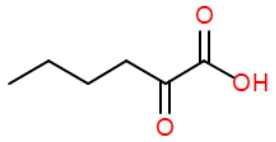 |
| 2-hydroxyadipic acid | -0.7 ± 1.7 (10 μM)<br>0.7 ± 1.0 | – | – | 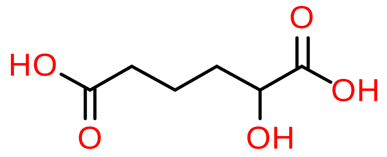 |
| 3-hydroxyadipic acid | 0.5 ± 2.7 (10 μM)<br>2.4 ± 1.6 | – | – | 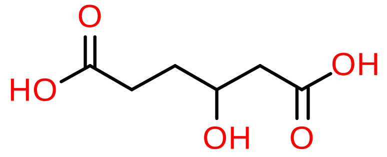 |
| omeprazole | 68.1 ± 0.4 | 102 ± 4 | n.d. | 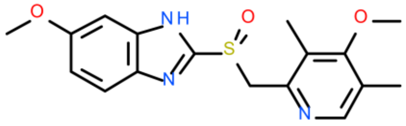 |
| pantoprazole | 52.8 ± 1.0 | 276 ± 13 | n.d. | 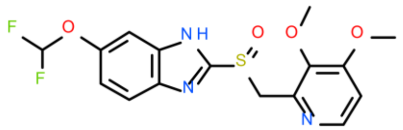 |
| esomeprazole | 62.9 ± 2.1 | 153 ± 4 | n.d. | 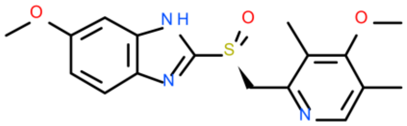 |
| tenatoprazole | 91.5 ± 1.8 | 27 ± 1 | 83 ± 34<br>(noncompetitive) | 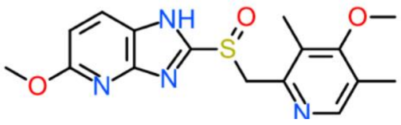 |

| | | | | |
|---|---|---|---|---|
| omeprazole sulfone | 44.4 ± 0.9 | 334 ± 6 | n.d. | 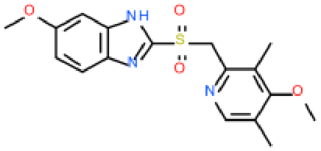 |

Activities were measured using 62.5 μM OP as substrate and the final concentration of DMSO was 1%, unless otherwise stated. [a]Inhibition relative to control (1% DMSO or water). [b]Compounds were tested at 300 μM, unless otherwise stated. [c]Compound dissolved in water. [d]n.d., not determined.

**Table S2. X-ray data collection and refinement statistics.**

| | DHTKD1:ThDP |
|---|---|
| **Data collection** | |
| Space Group | C 1 2 1 |
| Cell dimensions | |
| a,b,c (Å) | 332.314 72.6123 79.6752 |
| α, β, γ (°) | 90 91.8655 90 |
| Resolution (Å) | 49 - 2.1 (2.176 - 2.1) |
| $R_{sym}$ or $R_{merge}$ | 0.264 (2.031) |
| $R_{pim}$ | 0.098 (0.914) |
| $I/\sigma I$ | 5.9 (0.9) |
| CC(1/2) | 0.990 (0.313) |
| Completeness (%) | 99.9 (98.3) |
| Redundancy | 7.8 (5.8) |
| Average mosaicity | 0.22 |
| **Refinement** | |
| Resolution (Å) | 49-2.1 |
| No. reflections | 110798 (10904) |
| $R_{work}/R_{free}$ | 0.2138/0.2473 |
| No. atoms | |
| Protein | 13716 |
| Ligand/ion | 95 |
| Water | 1001 |
| B-factors | |
| Protein | 40.56 |
| Ligand/ion | 36.72 |
| Water | 41.05 |
| R.m.s deviations | |
| Bond lengths (Å) | 0.005 |
| Bond Angles (°) | 0.68 |

*Values in parenthesis are for highest-resolution shell. Data is from 2 crystals.

**Table S3. Oligonucleotides used for site-directed mutagenesis**.

| Primer | PAH cDNA position[a] | Sequence (5'→3')[b] |
|---|---|---|
| R163Q_F | 476-501 | CCACAGAAGAGC**A**AAAACATCTGTCG |
| R163Q_R | 476-501 | CGACAGATGTTTT**T**GCTCTTCTGTGG |
| L234G_F | 681-718 | GAATTTATTGACAGGCCTT**GG**GCAGTTCCCTCCAGAGC |
| L234G_R | 681-718 | GCTCTGGAGGGAACTGC**CC**AAGGCCTGTCAATAAATTC |
| Q305H_F | 897-931 | CAAAACTCGCGGCAGGCA**C**CAGTCTCGCCAAGACG |
| Q305H_R | 897-931 | CGTCTTGGCGAGACTG**G**TGCCTGCCGCGAGTTTTG |
| V360A_F | 1062-1099 | CAGAATTGGTGGGAGTG**C**GCATTTGATTGTTAATAACC |
| V360A_A | 1062-1099 | GGTTATTAACAATCAAATGC**G**CACTCCCACCAATTCTG |
| R455Q_F | 1347-1380 | CAAAATCATCAGAGCTC**A**AAAGAGCATTCCAGAC |
| R455Q_R | 1347-1380 | GTCTGGAATGCTCTTT**T**GAGCTCTGATGATTTTG |
| R715C_F | 2129-2159 | CCTGTCGAATAGAG**T**GTTTCCTGCAGATGTG |
| R715C_R | 2129-2159 | CACATCTGCAGGAAAC**A**CTCTATTCGACAGG |
| G729R_F | 2170-2202 | GAAGAGGGGGTGGAC**A**GAGACACTGTGAACATG |
| G729R_R | 2170-2202 | CATGTTCACAGTGTCTC**T**GTCCACCCCCTCTTC |
| P773L_F | 2301-2335 | GATGTTACTCAGGCTCC**T**GGCAGCCGTG TCAACTC |
| P773L_R | 2301-2335 | GAGTTGACACGGCTGCC**A**GGAGCCTGAGTAACATC |
| S777P_F | 2313-2347 | GCTCCCGGCAGCCGTG**C**CAACTCTTCAAGAAATGG |
| S777P_R | 2313-2347 | CCATTTCTTGAAGAGTTG**G**CACGGCTGCCGGGAGC |
| S862I_F | 2572-2601 | GATCATATTTGGA**T**TCAGGAGGAACCTCAG |
| S862I_R | 2572-2601 | CTGAGGTTCCTCCTGA**A**TCCAAATATGATC |

[a]GenBank accession number BC007955.2. [b]Mismatch nucleotides are shown in bold.

### References

1. T. Wagner, N. Barilone, P. M. Alzari, M. Bellinzoni, A dual conformation of the post-decarboxylation intermediate is associated with distinct enzyme states in mycobacterial KGD (alpha-ketoglutarate decarboxylase). *Biochem. J.* **457**, 425-434 (2014).

### NMR spectra

### $^1$H-NMR spectra of 1a.

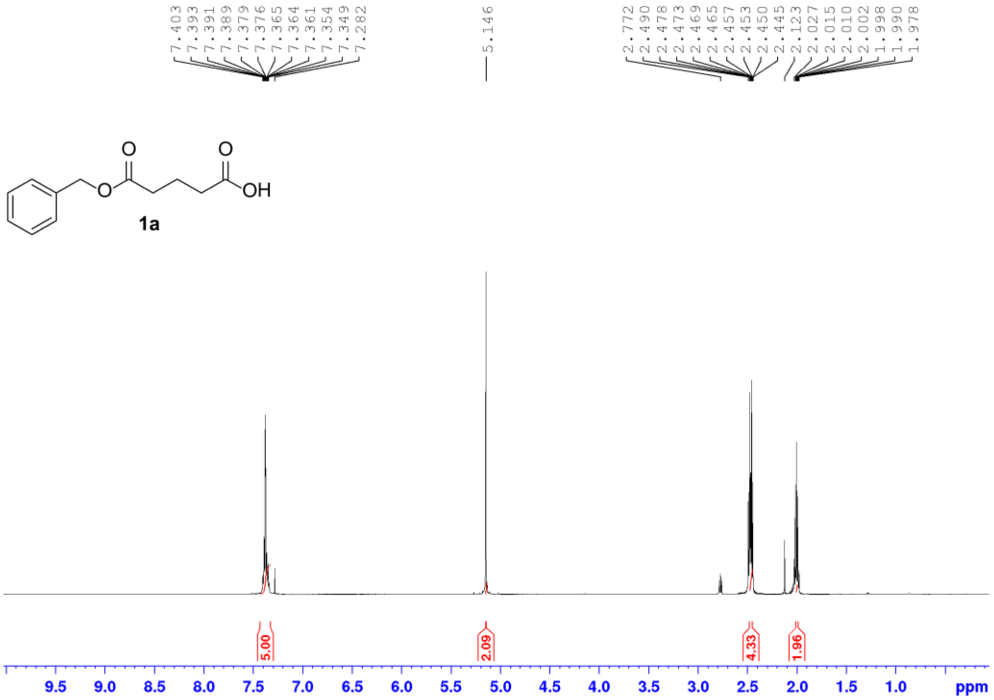

### $^1$H-NMR spectra of 2a.

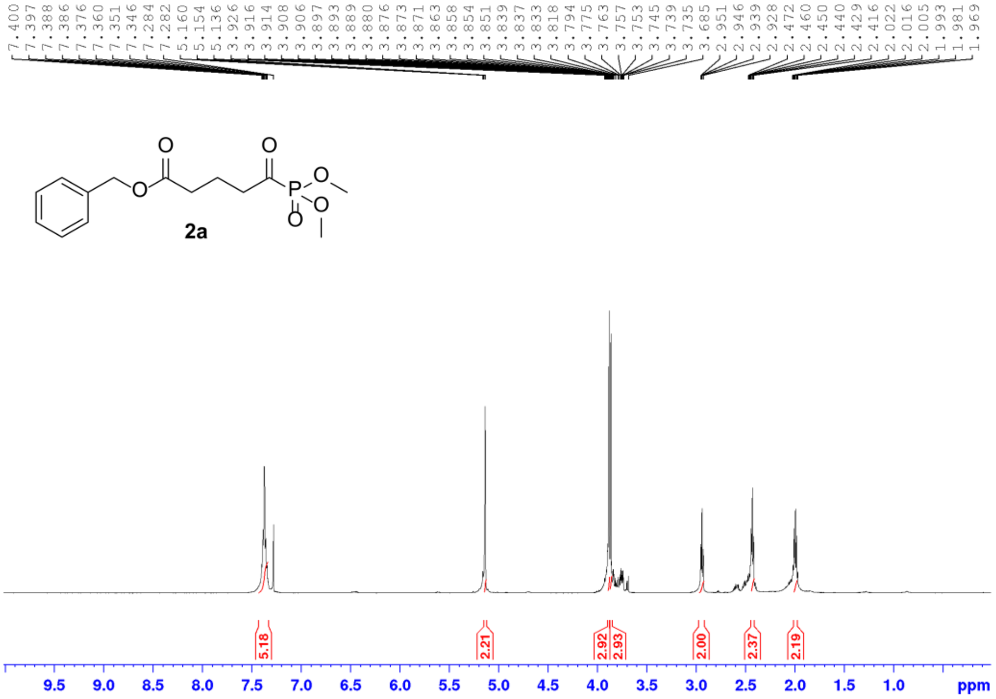

### 1H-NMR spectra of 3a.

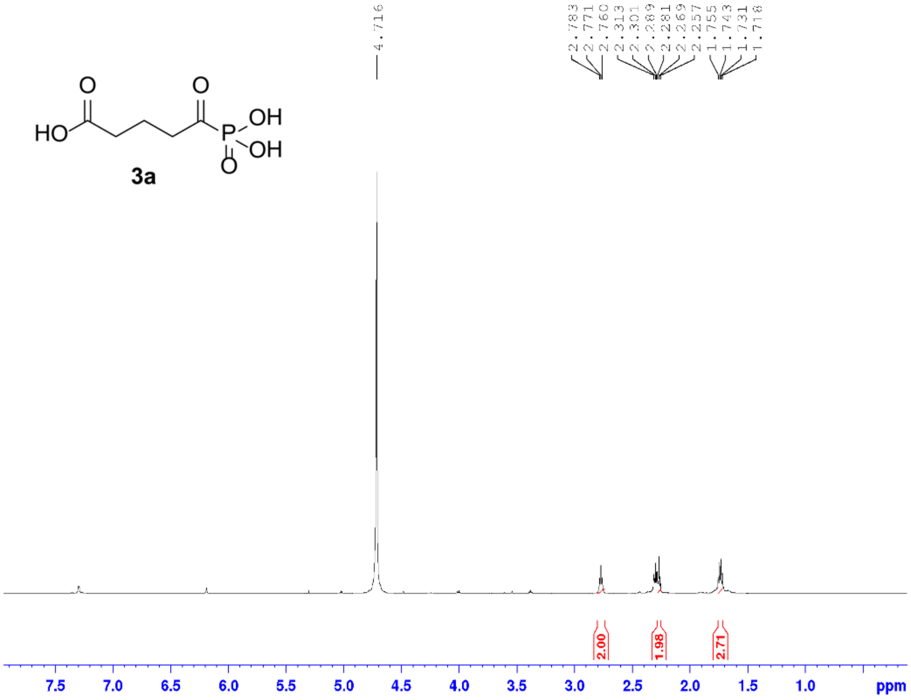

### 1H-NMR spectra of 1b.

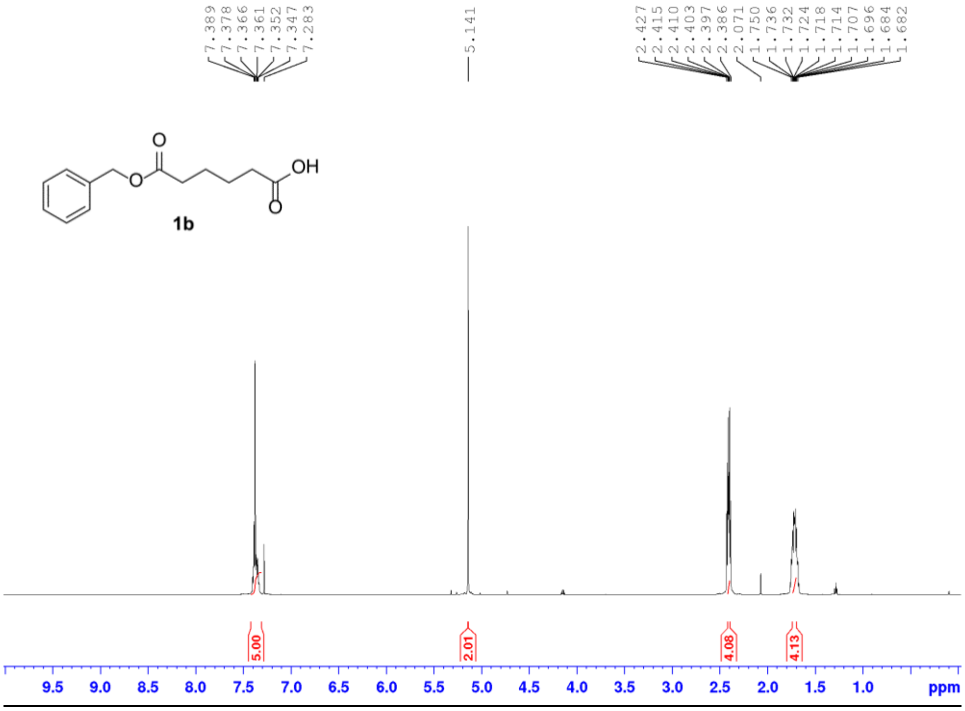

### [1]H-NMR spectra of 1bl.

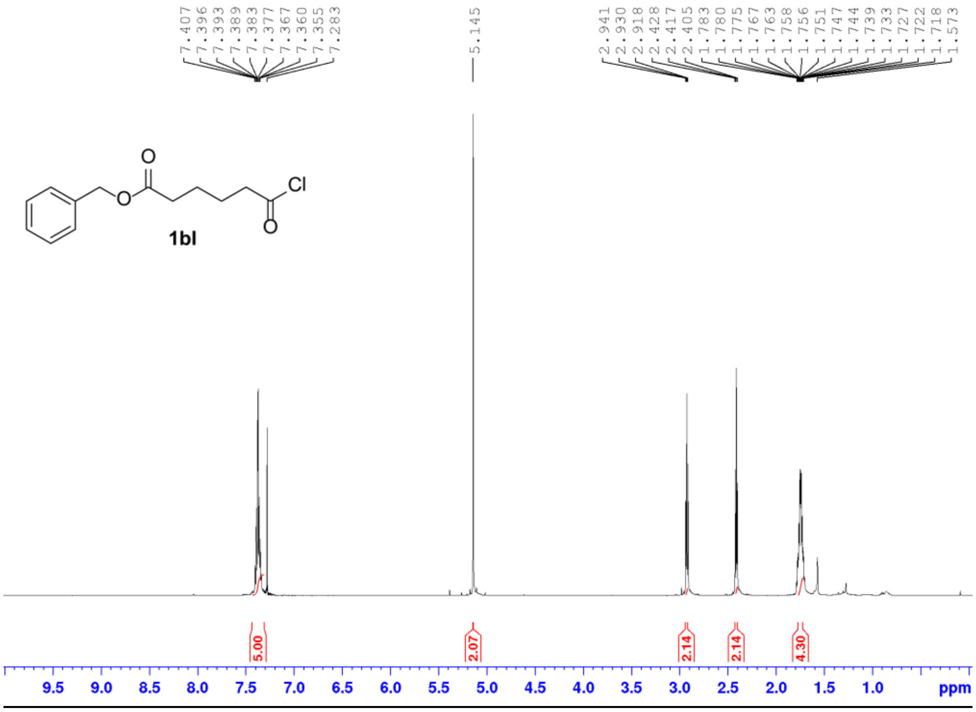

### [1]H-NMR spectra of 2b.

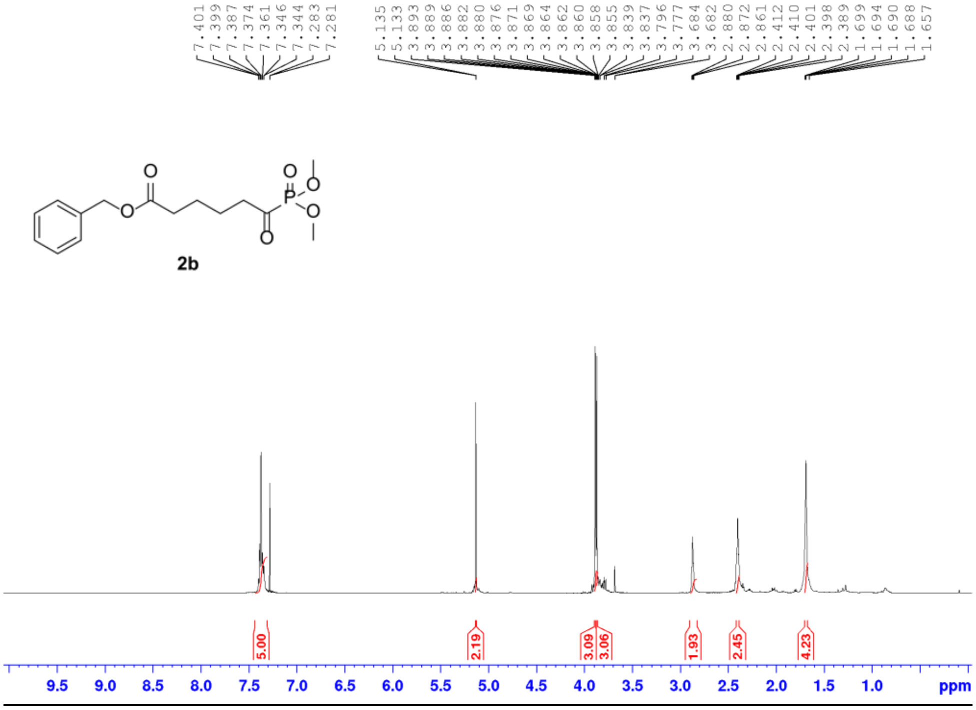

### [1]H-NMR spectra of 3b.

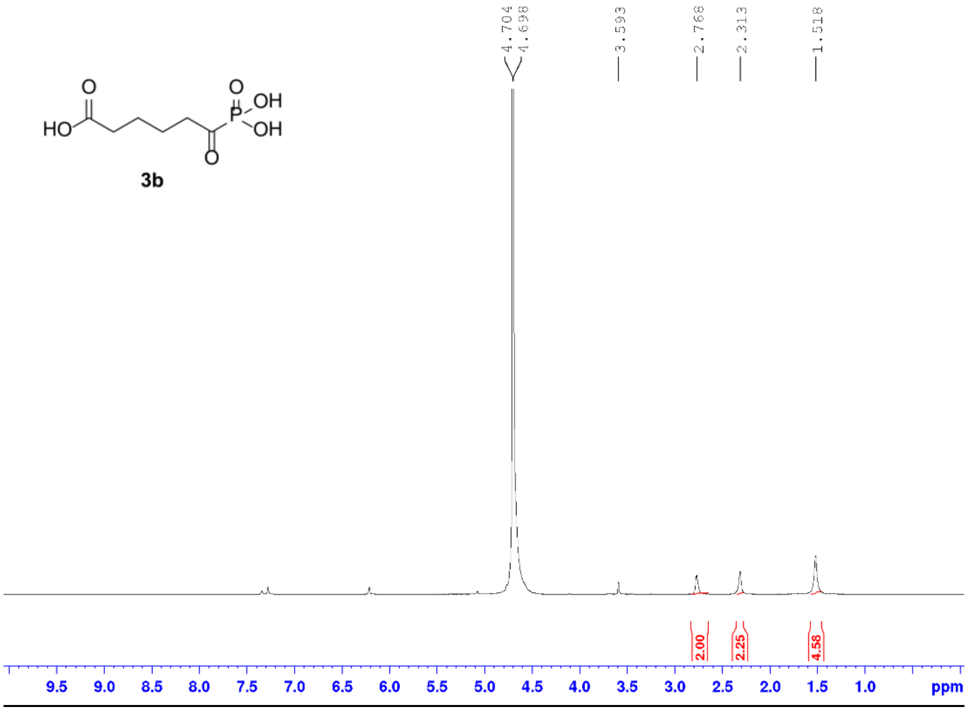

### [1]H-NMR spectra of 1c.

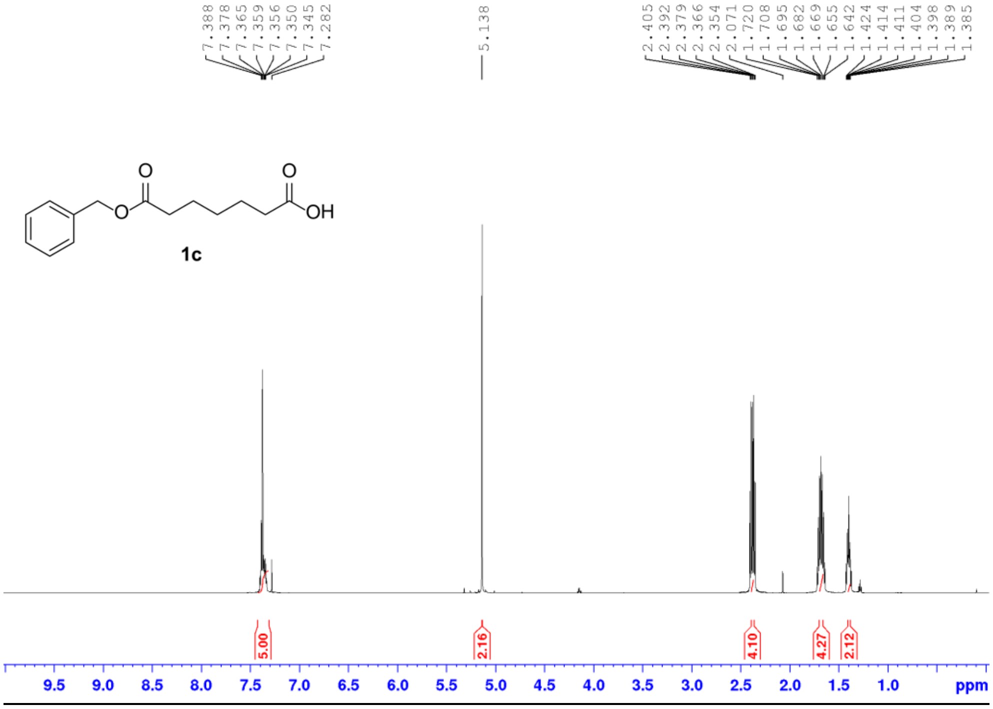

### [1]H-NMR spectra of 1cl.

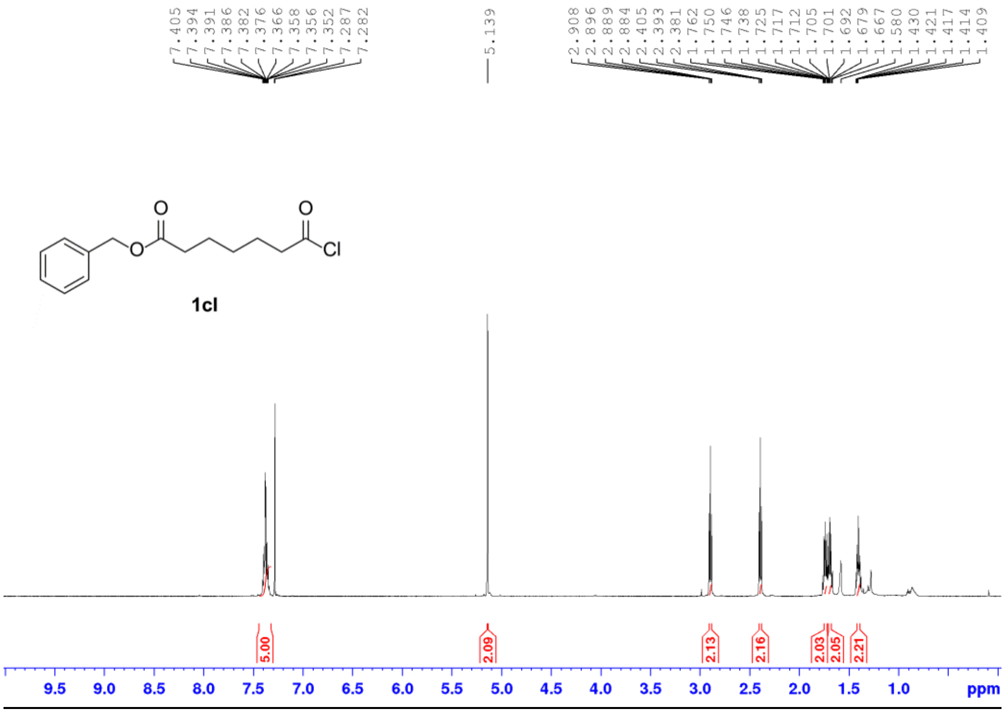

### [1]H-NMR spectra of 2c.

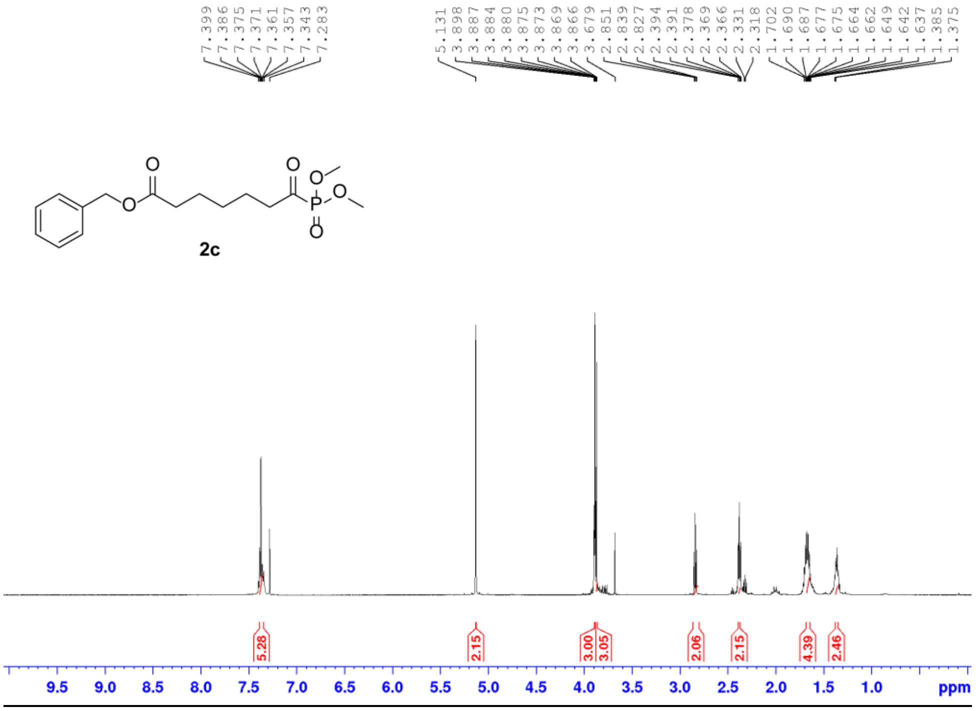

**[1]H-NMR spectra of 3c.**

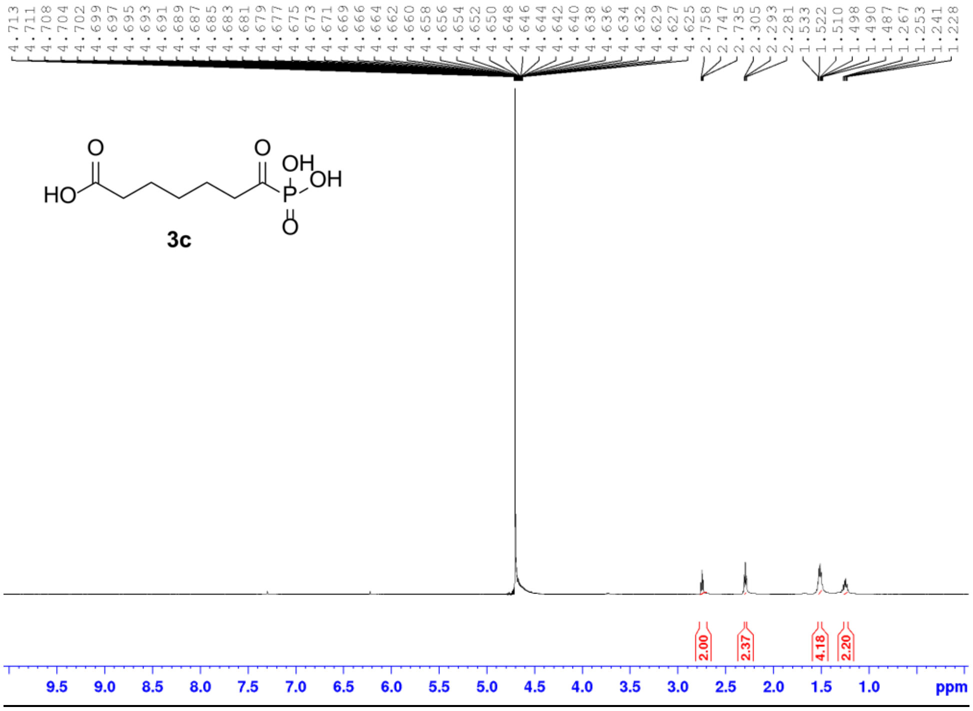
